# Working memory-based and -free reward prediction in a dual dopamine system in the basal ganglia

**DOI:** 10.1101/2023.03.06.531239

**Authors:** Tomohiko Yoshizawa, Yuuto Miyamura, Yuta Ochi, Riichiro Hira, Makoto Funahashi, Yutaka Sakai, Yilong Cui, Yoshikazu Isomura

## Abstract

Animals optimize their actions by predicting and verifying their outcomes (e.g., rewards). Reward prediction often requires working memory (WM)-based information. To elucidate the neural basis of WM-based reward prediction, we compared the activity of dopamine (DA) neurons in the ventral tegmental area (VTA) in an alternate reward condition (WM-based) with that in a random (WM-free) reward condition in rats and mice. Positron emission tomography revealed greater VTA activation in the WM-based than the WM-free condition. Lateral and medial VTA neurons displayed differential electrophysiological spike activities reflecting WM-based and WM-free reward prediction error, respectively. Furthermore, phasic DA release in the dorsal and ventral striatum changed as WM-based and WM-free classical conditioning progressed. Consistent with our WM-based model, reward acquisition caused a DA dip only in the dorsal striatum. Thus a dual DA system processes WM-based and WM-free reward prediction in parallel.

## Introduction

Humans and animals can optimize their behavior by predicting future outcomes (e.g., reward or punishment) of actions selected in accordance with the external environment. In classical and operant (in theory, reinforcement) learning experiments, the associations between cue, action, and outcome are formed through trial and error during trial repetition (Thorndike, 1911; Pavlov, 1927; Skinner, 1938). Such “contingency” formation requires some form of working memory (WM) and reference memory (RM). WM is temporary task-demanded memory valid for only one trial, while RM is a longer-term memory, consisting of general rules and external/internal states, that is commonly valid across trials (Olton and Samuelson, 1976).

A theoretical model of reinforcement learning explains the neural mechanism of reward prediction based on RM of past rewards. The learning depends on the prediction of future rewards according to the current sensory state and possible actions, i.e., “state value” and “action value,” respectively (Sutton and Barto, 1998). Functional brain imaging (O’Doherty et al., 2004; Tanaka et al., 2004) and neuronal activity recording (Samejima et al., 2005; Ito and Doya, 2009; Kim et al., 2009; Ito and Doya, 2015; Yoshizawa et al., 2018) studies have shown that reinforcement learning is implemented in the cortico-basal ganglia circuits. Dopaminergic neurons in the ventral tegmental area (VTA) and the substantia nigra pars compacta (SNc) encode the reward prediction error (RPE), defined as the discrepancy between actual and predicted rewards (Schultz et al., 1997). The striatal neurons, which receive the DA inputs from the midbrain VTA/SNc as well as glutamatergic inputs from the cerebral cortex, encode the state and action values of reinforcement learning (Samejima et al., 2005; Ito and Doya, 2009; Kim et al., 2009; Ito and Doya, 2015; Yoshizawa et al., 2018).

This learning sometimes utilizes a WM-based process as well. In WM-based reward prediction, forthcoming rewards can be directly predicted from the latest trials. For example, in a rock-paper-scissors game, if a person wins, they tend to continue the action in the next game, while if they lose, they are likely to switch to other actions (Wang et al., 2014). Such WM-based behaviors are referred to as “Win-Stay-Lose-Switch (WSLS).” We recently showed that dorsal striatum neurons encode previous action or reward when rats engaged in WSLS behavior in a choice task, but the neural coding was impaired when they failed to take the WSLS strategy due to insertion of a distractor into trials (Yoshizawa et al., 2022). Similarly, prolonging inter-trial intervals (ITIs) diminished WM-based WSLS behavior in mice (Iigaya et al., 2018). Moreover, several studies in humans reported that choice strategies were affected by altering the availability of WM by increasing the load of sensory information (Collins and Frank, 2012; Collins et al., 2014), adding another task in parallel (Otto et al., 2013a), or experiencing acute stress (Otto et al., 2013b).

These findings support the possibility that WM- and RM-based information may be adaptively processed according to task requirements. In fact, dopaminergic neurons encode different RPEs reflecting task structures in which a reward can be expected after a fixed number of no-reward trials (Nakahara et al., 2004) or after a time lapse (Starkweather et al., 2017). Such RPE coding disappears after lesioning of the orbitofrontal cortex (OFC) that reciprocally connects with the VTA (Takahashi et al., 2011), implying that computation of RPEs involves the cortico-basal ganglia circuits. However, it remains unknown how the midbrain dopamine (DA) system appropriately processes WM- and RM-based reward predictions to optimize future actions in reinforcement learning.

To address this issue, we compared the neural activity of the midbrain DA system in rodents when they were alternately rewarded and not rewarded (i.e., in a WM-based manner) versus randomly rewarded (i.e., in a WM-free manner) for correct trial performance. The alternating, but not random, reward condition enabled them to naturally predict the next outcome (reward or no reward) without any external instruction (Isomura et al., 2013). In the alternate- and random-reward task conditions, we employed (1) positron emission tomography (PET) to assess differentially activated brain areas, (2) electrophysiological spike recording from VTA neurons to evaluate the RPE signal, and (3) fluorescent fiber photometry to measure DA release in the dorsomedial striatum (DMS) and nucleus accumbens (NAc), which receive the midbrain DA inputs. It is noteworthy that we unexpectedly observed that acquisition of a reward (but not the absence of a reward) could cause a “dip” of DA release in a specific situation, which supports our WM-based and WM-free learning model in the dual midbrain DA system.

## Results

### Differentially activated brain areas in WM-based and WM-free reward prediction

Rats performed a lever-push task in either the alternate-reward (WM-based) or random-reward (WM-free) condition, immediately before they underwent 2-deoxy-2-[^18^F]fluoro-D-glucose PET (^18^F-FDG-PET) imaging. ^18^F-FDG was injected intravenously prior to the behavioral task performance on each PET scanning day. In each trial of the behavioral task, the ^18^F-FDG-injected rats continuously pushed a spout-lever (Kimura et al., 2012) on the wall of an operant chamber during a hold period. If the rats released the spout-lever in response to a go-cue tone, they were rewarded with a drop of water or received no reward according to the task condition (**Figure 1a**). In the alternate-reward condition, reward and no reward were alternately presented in every correct trial, whereas they appeared with random and equal probability in the random-reward condition. The rats were well trained to exactly predict a forthcoming reward in the alternate-reward condition, based on their WM information regarding the latest outcome. Therefore, we expected to detect which areas were more activated in WM-based vs. WM-free reward prediction by comparing PET signals in the whole brain between the alternate- and random-reward conditions.

**Figure 1.**
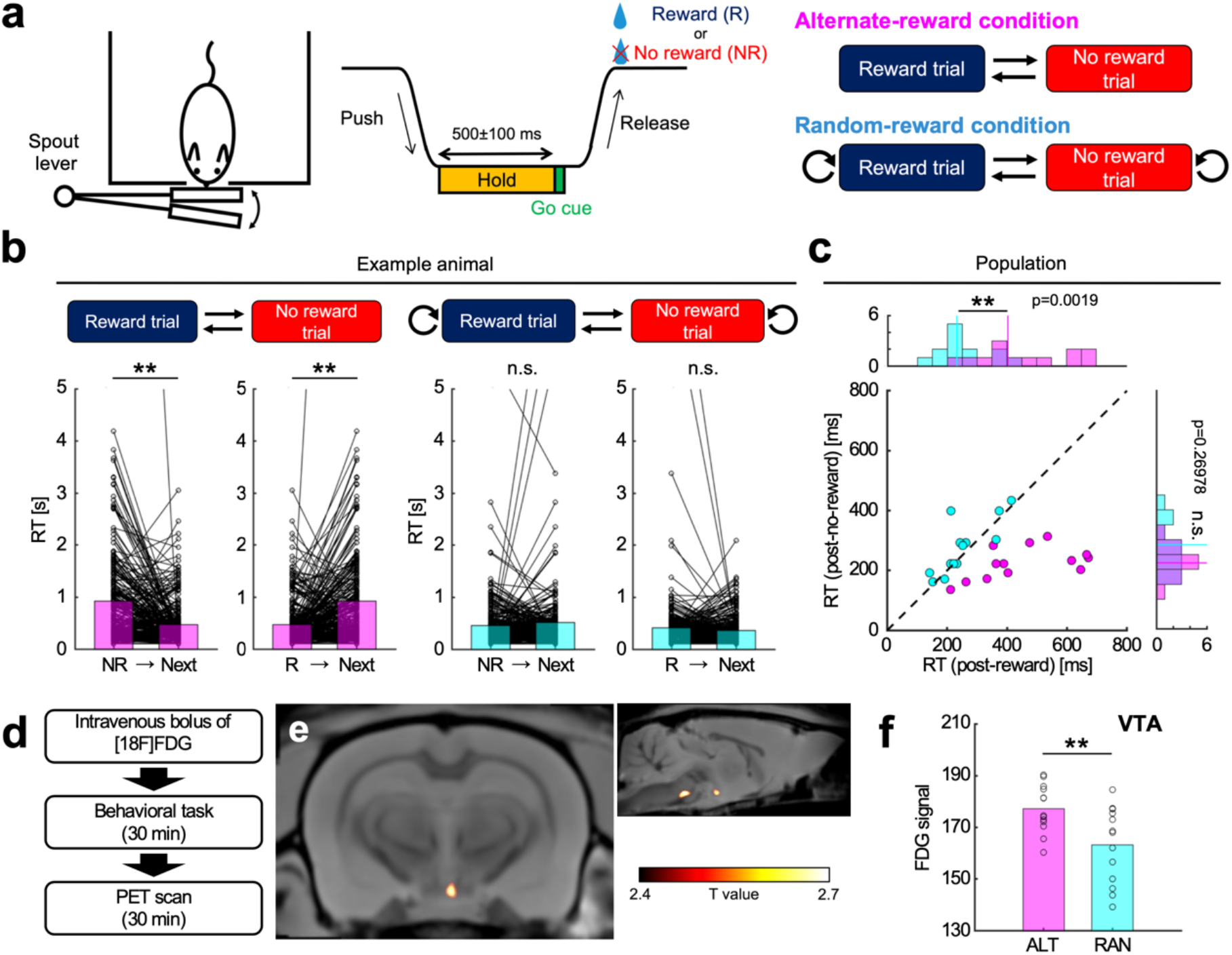
Positron emission tomography imaging revealed whole-brain activity during WM-based reward prediction. **(a)** Behavioral task. The experimental chamber was equipped with a spout-lever on a wall. After freely moving rats spontaneously pushed the spout-lever and maintained the pushing for 500±100 ms, a go-cue tone was presented. They received an outcome by releasing the spout-lever after the go-cue presentation. In the alternate-reward condition, they alternately performed a reward trial and a no-reward trial. In the random-reward condition, they randomly performed a reward trial and a no-reward trial with 50% probability. **(b)** Representative examples of rat performance. In the alternate-reward condition, the reaction time (RT) was shorter and longer in trials following no-reward and reward trials, respectively. In the random-reward condition, the previous outcome had no effect on the RT in the next trial. **: p < 0.01, n.s.: p ≥ 0.05, paired *t*-test. **(c)** Comparison of RT between conditions. The median RT in the post-reward trial was significantly longer in the alternate-reward condition than in the random-reward condition. In contrast, there was no significant difference in the post-no-reward trial. **p < 0.01, n.s.: p ≥ 0.05, Mann–Whitney *U* test. **(d)** Positron emission tomography (PET) scanning procedures. **(e)** Brain areas with increased ^18^F-FDG uptake in the alternate-reward condition compared to the random-reward condition. Increased ^18^F-FDG uptake was observed in the ventral tegmental area (VTA), and reticulotegmental nucleus of the pons. **(f)** ^18^F-FDG uptake in the VTA. The uptake was significantly greater in the alternate-reward condition than in the random-reward condition. ALT: alternate-reward condition, RAN: random-reward condition. **p < 0.01, unpaired *t*-test.

We compared the reaction time (RT) from the onset of the go cue to the lever release between the alternate- and random-reward condition groups. In a representative rat in the alternate-reward group, the RT was longer and shorter in subsequent trials following reward and no-reward trials, respectively (post-reward trials: p = 1.3e-07, post-no-reward trials: p = 3.0e-06, paired *t*-test, **Figure 1b**). Similar differences were observed in all the rats in the alternate-reward group. In the random-reward condition, however, the previous outcome did not affect the RT in the subsequent trials (post-reward trials: p = 0.90, post-no-reward trials: p = 0.91). Population analyses (alternate: 13 rats, random: 13 rats) showed that the RT in post-reward trials was significantly longer in the alternate-reward condition than in the random-reward condition (median RT; alternate: 403 ms, random: 232 ms, p = 0.0019, Mann– Whitney *U* test, **Figure 1c**). The RT in post-no-reward trials was not significantly different between the two conditions (alternate: 222 ms, random: 283 ms, p =0.27). The post-reward and post-no-reward trials were equivalent to no-reward and reward trials, respectively, in the alternate-reward condition. Therefore, these results indicated that in the alternate-reward condition, rats performed WM-based reward prediction using one-trial information on the latest outcome only.

After completing the behavioral session, the rats were immediately transferred to a PET scanner (**Figure 1d**). We obtained PET signals to identify brain areas with increased ^18^F-FDG uptake in the alternate- and random-reward groups (see Materials and Methods). Whole-brain analysis comparing the two groups revealed that in the alternate-reward group, ^18^F-FDG uptake was significantly increased in the primary auditory area (Au1), reticulotegmental nucleus of the pons (RtTg), pontine reticular nucleus, oral part (PnO) and caudal part (PnC), and VTA (p < 0.0125, uncorrected for primary survey, **Figure 1e**, Extended data Fig.1a). Among these areas, it is known that the VTA dopaminergic neurons encode RPE by spike activity (Cohen et al., 2012). Thus, we focused on the VTA for further analysis by setting a voxel of interest (VOI) on this region. VOI analysis confirmed that ^18^F-FDG uptake was significantly greater in the alternate-reward group than in the random-reward group (p = 0.0080, unpaired *t*-test, **Figure 1f**). Note that ^18^F-FDG uptake was not correlated with body weight, number of trials, or mean RT among sessions (Extended data Fig.1b, c, d, e).

We next computed functional connectivity based on the correlation of ^18^F-FDG uptake in the VTA with that in other brain areas to specify neural circuits for each of the alternate- and random-reward conditions. The functional connectivity of the VTA with the NAc, medial OFC, insular cortex, and thalamus waswas increased in the alternate-reward condition than in the random-reward condition (p < 0.005, Fisher’s Z-transformation test, **Figure 2a**). The functional connectivity of the VTA with the PnO, hypothalamus, mammillary body, amygdala, entorhinal cortex, and cerebellum was increased in the random-reward condition than in the alternate-reward condition (**Figure 2b**). In short, the functional connectivity between the VTA and rostral brain areas was strengthened in the alternate-reward condition, while that between the VTA and caudal brain areas was strengthened in the random-reward condition (**Figure 2c**). To further analyze the functional coupling between the VTA and NAc, VOI analysis was applied to these areas. In the random-reward condition, ^18^F-FDG uptake in the VTA was significantly negatively correlated with that in the NAc (r = -0.78, p = 0.0017, Pearson correlation analysis, **Figure 2d**). In contrast, there was no significant correlation in the alternate-reward condition (r = 0.33, p = 0.27). The correlation coefficients were significantly different between the alternate- and random-reward conditions (p = 0.0019, Fisher’s Z-transformation test). These results suggest that the VTA-NAc pathway contributed to WM-free reward prediction.

**Figure 2.**
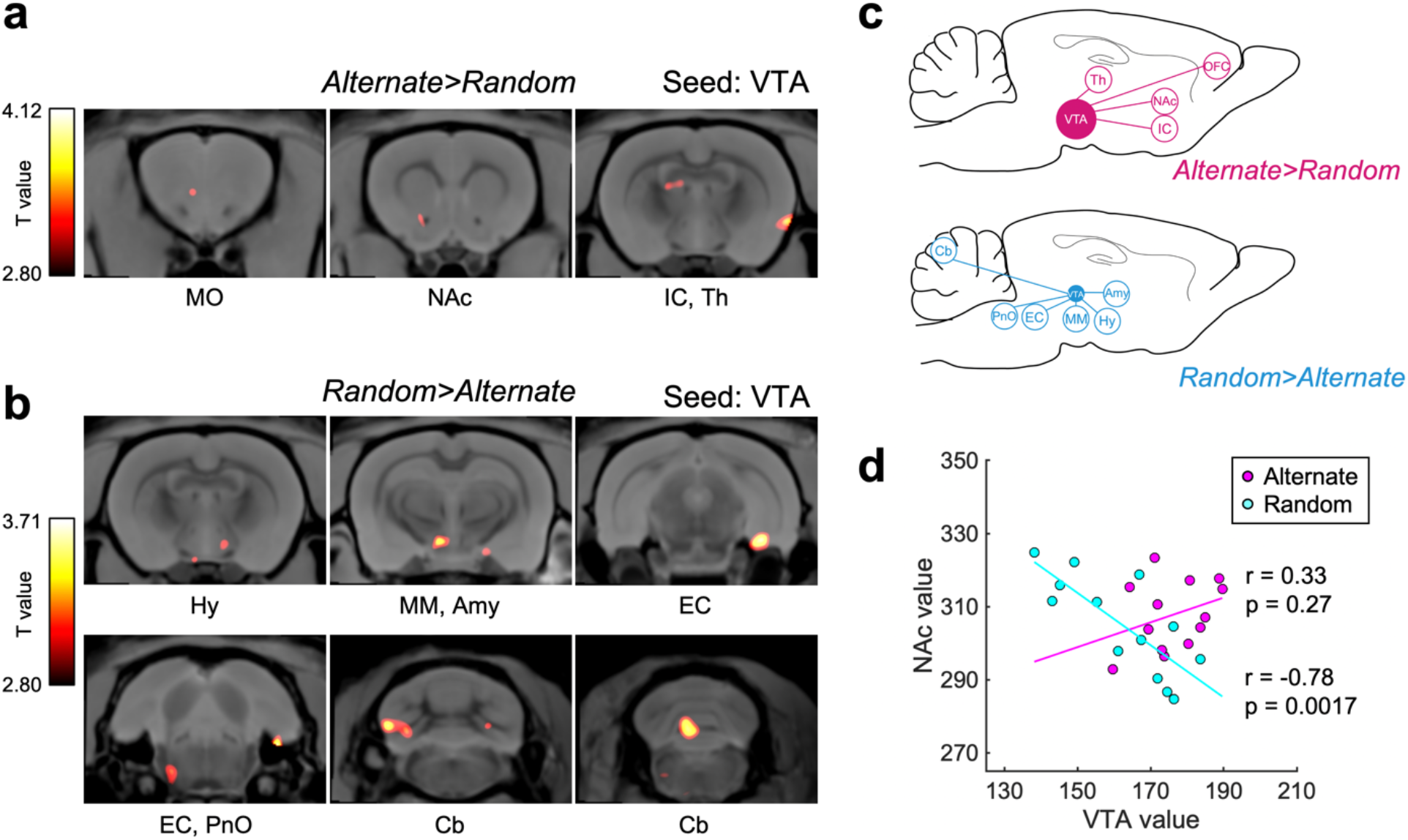
Distinct neural circuits contributed to responses in the alternate- and random-reward conditions. **(a)** Brain areas with increased functional connectivity (FC) to the VTA in the alternate-reward condition. MO: medial orbitofrontal cortex, IC: insular cortex, NAc: nucleus accumbens, Th: Thalamus. **(b)** Brain areas with increased FC to the VTA in the random-reward condition. Hy: Hypothalamus, MM: Mammillary body, Amy: Amygdala, EC: Entorhinal cortex, PnO: Pontine reticular nucleus, oral part, Cb: Cerebellum. **(c)** Schematic representation of the results of FC analysis. VTA activities tended to correlate with activities of anterior and posterior brain areas in the alternate- and random-reward conditions, respectively. **(d)** Functional coupling between the VTA and NAc. Significant negative functional coupling was observed in the random-reward condition. Magenta and cyan lines indicate regression lines. Pearson correlation analysis.

### Differential activity of VTA neurons in WM-based and WM-free reward prediction

To investigate the difference in single-neuron activity between the two reward conditions, an electrophysiological spike recording technique was applied to VTA neurons while identical rats performed trials in both reward conditions under head fixation. In each trial, rats pushed and held a spout-lever until the presentation of a go-cue tone. If the rats pulled the spout-lever in response to the go-cue, they received a reward or no reward after a delay period (**Figure 3a, b**). We designed three types of trial blocks in which the alternate-reward, random-reward, and 100%-reward conditions were included in one task session (**Figure 3c**). In the alternate-reward condition, correct push-hold-pull actions were alternately rewarded and not rewarded. In the random-reward condition, the same actions were randomly rewarded with a 50% probability. In the 100%-reward condition, rats always received a reward for every correct action. Thus, WM-based reward prediction using information on the latest outcomes was possible only in the alternate-reward condition, while the net likelihood of expected rewards was equal (50%) between the alternate- and random-reward conditions. In the 100%-reward condition, the state and action values of go-cue signal and their action were twice as great as in the other two conditions. An important point is that unlike the above PET experiments, the same individuals were subjected to these three conditions for comparison in each session. Thereby, we expected to evaluate the contribution of single VTA neurons to WM-based or WM-free reward prediction.

**Figure 3.**
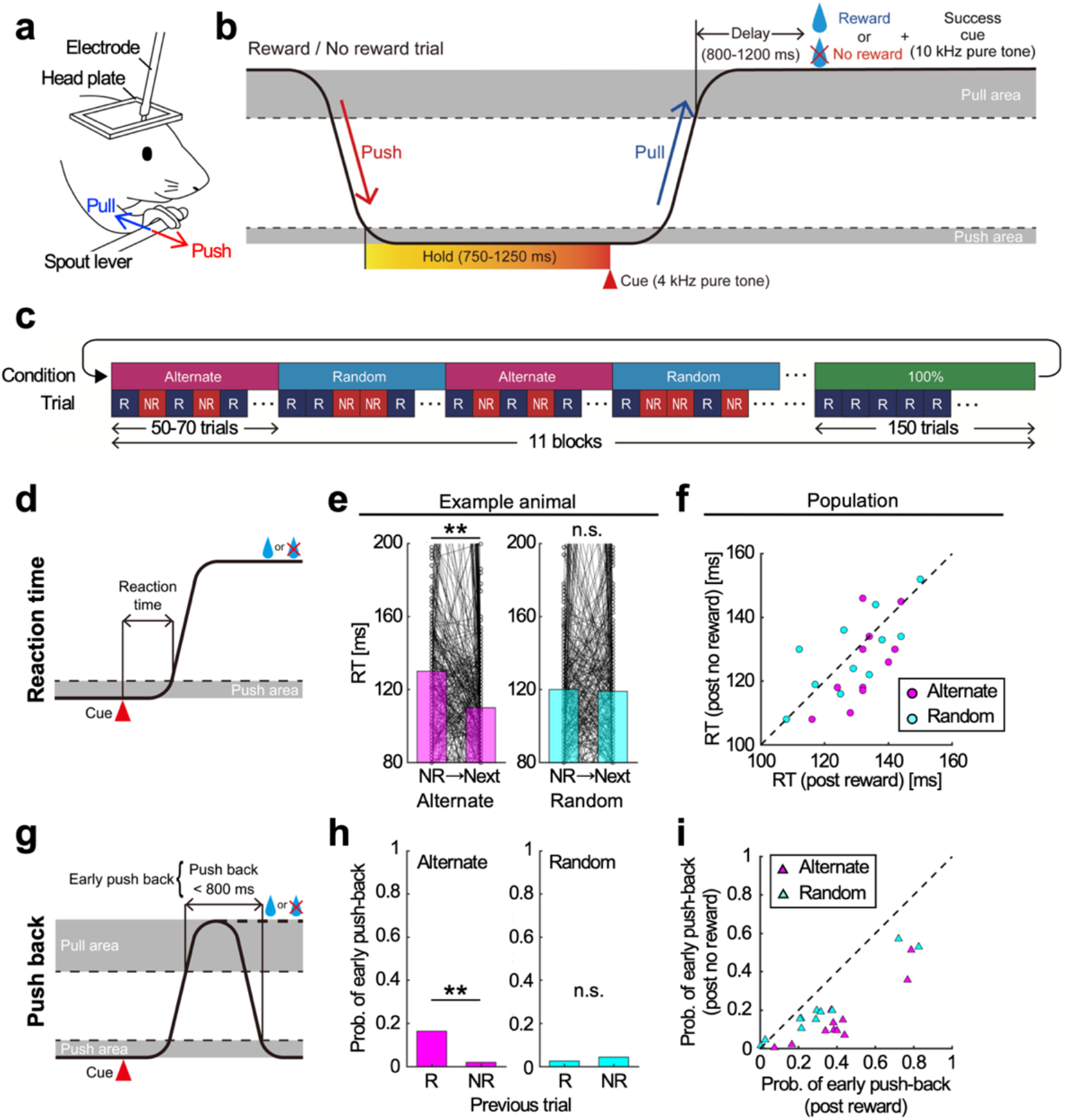
Rat behavior involving pushing or pulling the experimental lever reflected WM-based reward prediction. **(a)** Schematic illustration of the behavioral apparatus. The head and body of each rat was restrained by a metal frame and tube. They pushed and pulled a spout-lever with their right forelimb. **(b)** Time course of a lever push-pull task. A go-cue tone was presented when the rats continuously pushed the spout-lever for 750–1250 ms. When they pulled the spout-lever after the go-cue presentation, they received an outcome after a short delay. **(c)** Switching schedule of task conditions in a session. The task consisted of three reward conditions: one in which a lever push-pull behavior was alternately rewarded (alternate-reward condition), another in which the same operant behavior was randomly rewarded with a 50% probability (random-reward condition), and the last in which the same operant behavior was always rewarded (100%-reward condition). The alternate- and random-reward conditions consisted of 50–70 trials. The 100%-reward condition consisted of 150 trials. The 100%-reward condition appeared every 10 times the alternate- and random-reward condition were switched. The alternate-, random-, and 100%-reward conditions were switched without any external cue. **(d)** Definition of RT. RT was the time from go-cue onset to lever-pull onset. **(e)** A representative example of RT in a single session. In the alternate-reward condition, RTs were shorter and longer in the trials following no-reward and reward trials, respectively. In the random-reward condition, the previous outcome had no effect on the RT in the next trial. **: p < 0.01, n.s.: p ≥ 0.05, paired *t*-test. **(f)** RTs in post-reward and post-no-reward trials. In the alternate-reward condition, the RT was significantly shorter in the post-no-reward trial than in the post-reward trial. There was no significant difference in the random-reward condition. **(g)** Definition of early push-back. Early push-back behavior was an action that returned the spout-lever to the push position within 800 ms after go-cue presentation. **(h)** A representative example of the probability of fast push-back in a session. In the alternate-reward condition, the probability was higher in the post-reward trial than in the post-no-reward trial. In the random-reward condition, there was no significant difference. **: p < 0.01, n.s.: p ≥ 0.05, Chi-squared test. **(i)** Probabilities of early push-back in post-reward and post-no-reward trials. In both conditions, the probability was significantly higher in the post-reward trial than in the post-no-reward trial.

We compared RTs between the alternate- and random-reward conditions in each session (**Figure 3d**). In a representative rat, RTs were shorter in trials following no-reward trials in the alternate-reward condition (p = 1.5e-09, paired *t*-test, **Figure 3e**). In the random-reward condition, the latest outcome did not affect the RT in the next trial (p = 0.39). Population analyses of all 11 sessions revealed that the RT in the alternate-reward condition was significantly shorter in post-no-reward trials than in post-reward trials (median RT; post-reward: 132 ms, post-no-reward: 126 ms, p = 0.049, Wilcoxon signed-rank test, **Figure 3f**), whereas the RT in the random-reward condition was not significantly different between them (post-reward: 129 ms, post-no-reward: 130 ms, p = 0.86). These results in head-fixed rats were consistent with those observed in freely-moving rats in the PET experiments.

We also counted the number of times that rats exhibited early push-back behavior, defined as returning the spout-lever into the push position during the delay period before the possible outcome (**Figure 3g**). Early push-back behavior was considered a predictive behavior of the no-reward outcome, because if rats expected a reward, they needed to hold the spout-lever near their mouth in the pull position rather than push it back (**Figure 3a**). In the alternate-reward condition, an example rat exhibited a higher probability of early push-back in post-reward trials than in post-no-reward trials (post-reward: 0.16, post-no-reward: 0.020, p = 9.2e-10, chi-squared test, **Figure 3h**), whereas there was no significant difference in the random-reward condition (post-reward: 0.026, post-no-reward: 0.044, p = 0.23). Population analysis also revealed that the probability of early push-back in the alternate-reward condition was significantly higher in post-reward trials than in post-no-reward trials (median probability; post-reward: 0.38, post-no-reward: 0.095, p = 9.8e-04, Wilcoxon signed-rank test, **Figure 3i**). These results indicate that rats conducted WM-based reward prediction in the alternate-reward condition.

We performed electrophysiological spike recording by stereotaxically inserting a multichannel silicon probe into the VTA of rats performing the lever-pull task (**Figure 4a, b, c**). We isolated 165 VTA neurons from four rats. These neurons were further classified into putative dopaminergic neurons and GABAergic interneurons according to a clear bimodal distribution of their spike duration (**Figure 4d**). Putative GABAergic neurons showed a significantly higher basal firing rate (1.27 ± 0.78 Hz, n = 20 neurons) than putative dopaminergic neurons (0.72 ± 0.062 Hz, n = 145 neurons, p = 0.037, one-tailed Mann–Whitney *U* test), consistent with previous reports (Cohen et al., 2012; Mohebi et al., 2019). To analyze reward-related neurons, we first compared the number of spikes 300 ms before and after the onset of reward delivery. Of the 145 putative dopaminergic neurons, 61% (89/145 neurons) showed significantly positive responses to rewards in at least one of the alternate-, random- and 100%-reward conditions (p < 0.05/3, paired *t*-test followed by Bonferroni correction). No neurons showed significantly negative responses to rewards. Twenty-seven percent (24/89 neurons) of the reward-responsive neurons had significantly different responses to rewards between at least one pair of the three reward conditions (p < 0.05/3, unpaired *t*-test followed by Bonferroni correction). Of these, 71% (17/24) showed significantly different reward responses between the random- and 100%-reward conditions. In particular, 88% (15/17) of these neurons were so-called RPE neurons that showed significantly stronger reward responses in the random-reward condition than in the 100%-reward condition (Extended Data Fig.2). Also, 50% (12/24) and 17% (4/24) displayed different activity between the alternate- and 100%-reward conditions and between the alternate- and random-reward conditions, respectively.

**Figure 4.**
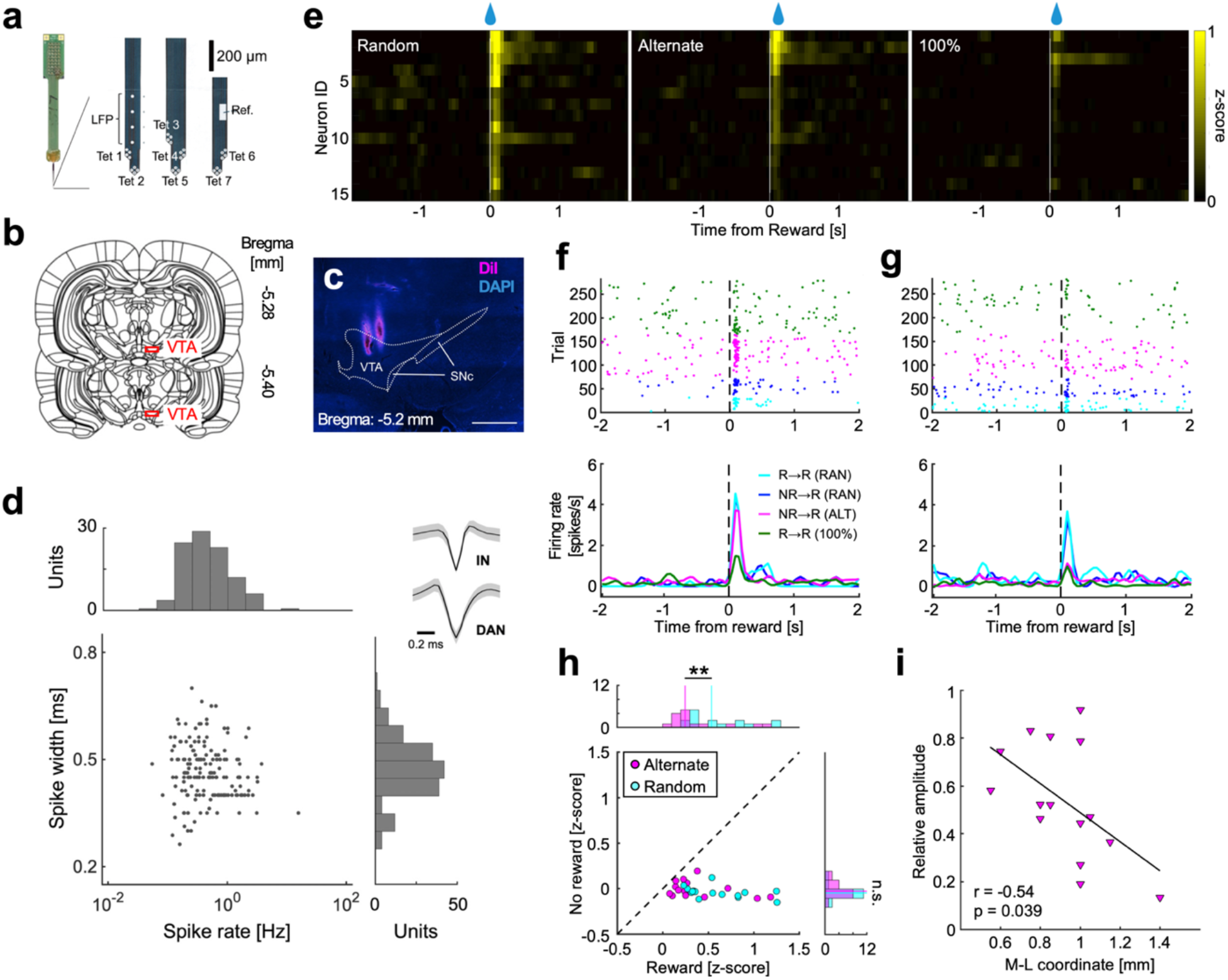
VTA neural activity encoded WM-based reward prediction error. **(a)** 32-ch silicone probe. The probe had three shanks, with two sets of tetrodes placed on the shanks at each end and three sets on the central shank. **(b)** VTA recording sites. The electrode was inserted into the VTA (red squares) before the start of a session and then removed after the end of the session. The recording sites were slightly changed in each recording session. Each diagram represents a coronal section referenced to the bregma (Paxions and Watson, 1998). **(c)** DAPI-stained coronal section showing the recording sites in the VTA. Electrode tracks were visualized with fluorescent DiI. Scale bar: 200 µm. **(d)** Classification of putative VTA dopaminergic neurons (DAN). The putative DAN were distinguished from the putative GABAergic interneurons (IN) by their biphasic spike width distribution. The insets show representative average spike waveforms of putative IN and DAN. **(e)** Normalized activity patterns of all reward prediction error (RPE) neurons. Indexes of neurons were sorted based on the amplitude of the reward response under the alternate-reward condition. **(f)** Raster plots and peri-event time histograms (PETHs) of a representative WM-free RPE neuron. The amplitude of the reward response did not differ between in the alternate- and random-reward conditions. The amplitude was smaller in the 100%-reward condition than in the other conditions. **(g)** Raster plots and PETHs of a representative WM-based RPE neuron. The Amplitude of the reward response was smaller in the alternate-reward condition than in the random-reward condition. In addition, the amplitude was similar between in the alternate-reward condition and the 100% reward condition. **(h)** Amplitude of the spike response to reward and no reward in the alternate- and random-reward conditions. The amplitude of the response to reward was significantly smaller in the alternate-reward condition than in the random-reward condition, whereas the response to no reward was not significantly different. **: p < 0.01, n.s.: p ≥ 0.05, Mann–Whitney *U* test. **(i)** Relation between recording position and response to the alternate reward. To quantify RPEs occurring in the alternate-reward condition, we normalized values so that the amplitude of the response to the random reward was 1 and that of the response to the 100% reward was 0, then obtained the relative amplitude of the response to the alternate reward. The relative amplitude of each RPE neuron was negatively correlated with the medial-lateral (M-L) coordinate of the recording position (r = -0.54, p = 0.039, Pearson correlation analysis). Black line indicates a regression line.

To further examine the reward response of all 15 RPE neurons, z-scored peri-event time histograms (PETHs) of their spike activity were arranged in the order of the reward response amplitude in the alternate-reward condition (**Figure 4e**). The reward responses for the alternate-reward condition ranged between those in the random-reward condition (larger) and the 100%- condition (smaller). A representative neuron responded most strongly to random rewards, regardless of previous outcomes, and to alternate rewards as much as the random rewards (**Figure 4f**). The same neuron showed only a weak response in the 100%-reward condition. Another neuron also responded strongly to random rewards, but it responded only weakly to alternate rewards and 100% rewards (**Figure 4g**). The former corresponds to a large RPE reflecting WM-free reward prediction, whereas the latter corresponds to a small RPE reflecting WM-based reward prediction in the alternate-reward condition. Population analysis of the 15 RPE neurons revealed that their reward responses were significantly lower in the alternate-than in the random-reward condition (median z-score; alternate: 0.25, random: 0.54, p = 0.014, Mann–Whitney *U* test, **Figure 4h**), whereas the no-reward responses were not significantly different between these two conditions (alternate: -0.022, random: -0.042, p = 0.30). To quantify the relative degree of RPEs occurring in the alternate-reward condition, we normalized them by defining the 100%-reward response and random-reward response to be 0 and 1, respectively; e.g., 0.92 and 0.19 for the neurons in **Figure 4f and 4g**, respectively. These RPE neurons displayed different relative RPEs in the alternate-reward condition depending on the medial-lateral (M-L) coordinate of the recording position in the VTA (r = -0.54, p = 0.039, Pearson correlation analysis, **Figure 4i**). Relative RPEs did not significantly correlate with the anterior-posterior (A-P) or dorsal-ventral (D-V) coordinate (Extended Data Fig.3). These results indicate that medial and lateral VTA neurons tended to encode WM-free and WM-based RPEs, respectively.

### Differential DA release in WM-based and WM-free reward prediction as learning progresses

We found spatially different RPE distribution according to the M-L width of the VTA (0.6∼1.4 mm lateral in rats; see **Figure 4i**). Anatomical investigations indicated that the medial half of our recorded sites (M-L 0.6∼1.0 mm) corresponded to the parabrachial pigmented nucleus of the VTA, which provides NAc-projecting dopaminergic neurons, whereas the lateral half (M-L 1.0∼1.4 mm) was the lateral VTA and part of the SNc, which provide DMS- as well as NAc-projecting dopaminergic neurons (Hilário and Costa, 2008; Parker et al., 2016; Saunders et al., 2018). A recent study reported that dopaminergic neurons in the medial and lateral VTA displayed RPE- and salience-related spike activities, respectively, in classically conditioned mice (Cai et al., 2020). Taking these observations and our own into account, the question arises as to how DA release into the DMS and NAc changes as outcome learning progresses. To address this question, we performed fiber photometry experiments to measure phasic DA release in the DMS and NAc at different stages of classical learning.

In the PET and electrophysiology experiments, it was more advantageous to use rats than mice in order to precisely distinguish the brain areas in terms of size. On the other hand, we had sufficient previous experience showing that head-fixed mice learned spout-licking behavior in a classical conditioning task in a few days (Yoshizawa et al., 2018), which made it possible to perform DA measurement across sessions. Hence, we trained mice to perform a classical conditioning task under head fixation for DA measurement by fiber photometry (**Figure 5a**). The classical conditioning consisted of a conditioned stimulus (a cue tone for 2 s) and an unconditioned stimulus (outcome: a drop of water as a reward or none) in the alternate- or random-reward condition (**Figure 5b**). The first fiber photometry session started after the establishment of reward-predictive licking behavior in naïve mice prior to an actual reward in the alternate-reward condition. Three sessions in the alternate-reward condition (ALT1 to ALT3) were followed by three sessions in the random-reward condition (RAN1 to RAN3) (**Figure 5c**).

**Figure 5.**
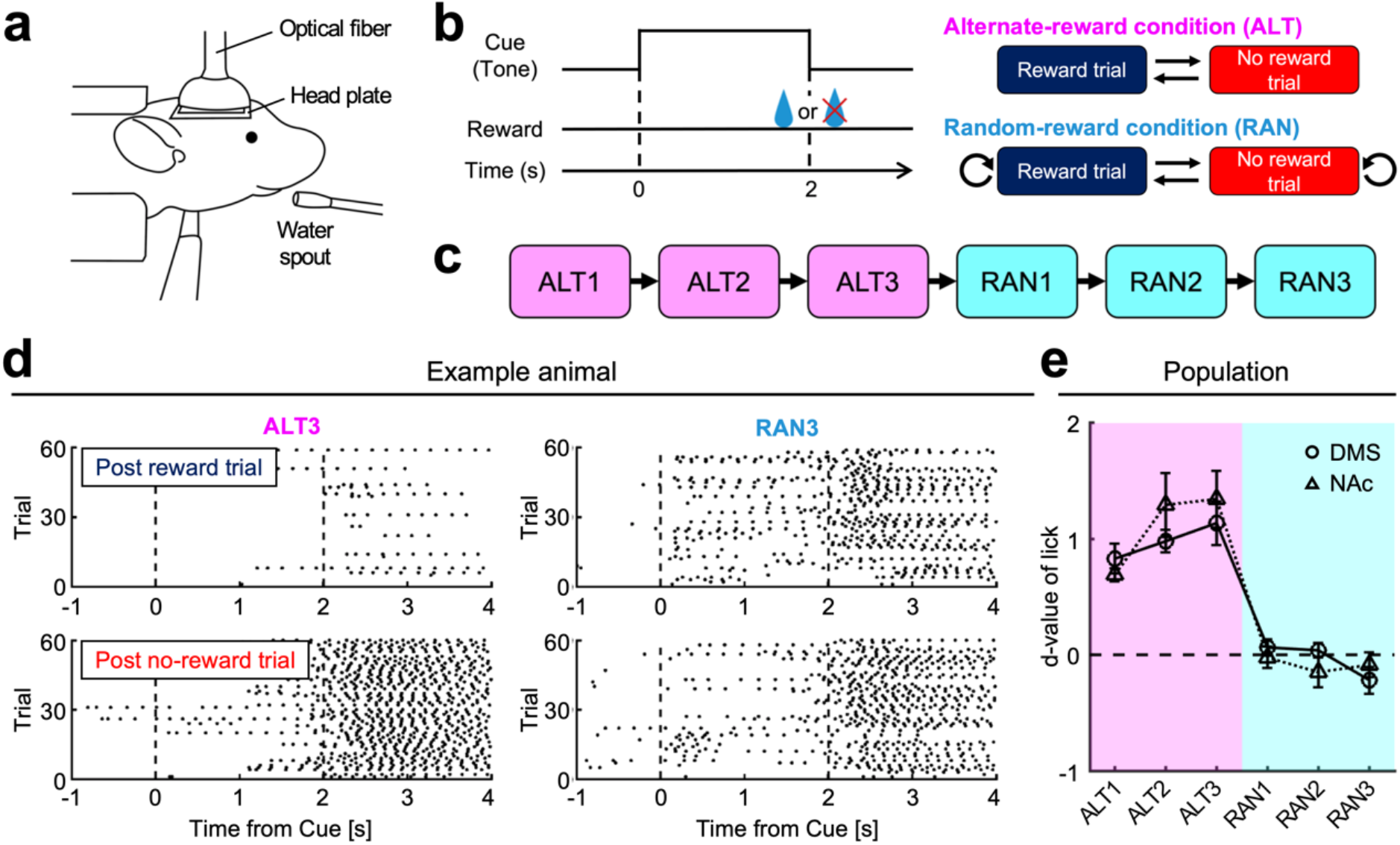
Mice showed WM-based reward-predictive licking behavior. **(a)** Schematic illustration of the behavioral apparatus. The head and body of each mouse was restrained by a metal frame and tube. A water spout was placed in front of its mouth. Spout-licking behavior was monitored by an infrared sensor. The optical fiber was inserted into the brain. **(b)** Time course of a classical conditioning task. In each trial, a cue tone was presented for 2 s, followed by an outcome. In the alternate-reward condition, a drop of water was alternately delivered or not delivered as a reward. In the random-reward condition, it was delivered randomly with 50% probability. **(c)** Task schedule. Three successive sessions of the alternate-reward condition (ALT1 to ALT3) were followed by three sessions of the random-reward condition (RAN1 to RAN3). **(d)** Examples of reward-predictive spout-licking behavior in ALT3 and RAN3. In ALT3, the post-no-reward trial was a reward trial. The reward-predictive licking behavior was observed during cue presentation in the post-no-reward trial. On the other hand, the post-reward trial was a no-reward trial. The reward-predictive licking behavior was not observed. In RAN3, the reward-predictive licking behavior was observed in both post-reward and post-no-reward trials. Black dots indicate spout-licking behaviors. **(e)** Effect size of task condition. Effect size *d* was the normalized difference of the reward-predictive licking behavior between the post-no-reward trial and the post-reward trial. The magnitude of *d* was significantly larger in the alternate-reward condition than in the random-reward condition.

In the alternate-reward sessions, reward-predictive licking by a representative mouse was more frequently observed in post-no-reward trials than post-reward trials (number of licks in 0.5 s before the outcome; post-reward: 0.14 ± 0.078, post-no-reward: 1.2 ± 0.16, p = 4.1e-08, unpaired *t*-test, **Figure 5d**), whereas there was no significant difference in the random-reward sessions (post-reward: 0.80 ± 0.14, post-no-reward: 0.72 ± 0.14, p = 0.69). One-way ANOVA revealed a significant main effect of the alternate-reward sessions (p = 0.031, **Figure 5e**) on the difference in reward-predictive licking between post-no-reward and post-reward trials (d-value; see Materials and Methods). Post hoc comparison using Tukey’s honestly significant difference test in the alternate-reward session revealed that the d-value was significantly larger in ALT3 than in ALT1 (ALT1: 0.77 ± 0.073, ALT3: 1.2 ± 0.15, p = 0.031), indicating learning progress. Subsequently, the d-values decreased to near zero in RAN1-3, demonstrating that reward-predictive licking was not specific to post-no-reward trials. Moreover, the longer the ITI was extended, the smaller the d-value became in four other mice under the alternate-reward condition (Extended data Fig.4). When the ITI was over 60 s, reward-predictive licking was observed with the same frequency in both post-reward and post-no-reward trials. These results suggested that the mice conducted WM-based reward prediction in the alternate-reward condition of classical conditioning.

Using fiber photometry with the fluorescent dopamine sensor dLight1.1, we evaluated DA release dynamics in the DMS and NAc of mice classically conditioned through these six sessions (Patriarchi et al., 2018) (**Figure 6a**, Extended Data Fig.5). In the DMS of a representative mouse, the DA signal (z-scored dLight1.1 fluorescence) was transiently elevated at the time of cue presentation (**Figure 6b**). The correlation between the trial order and cue responses was not significant through the alternate-reward sessions (correlation coefficient r_cue_ = 0.044, p = 0.55, Pearson correlation analysis), indicating that the cue response had no learning effect on DA release in the DMS (**Figure 6c**). The second elevation of the DA signal was observed when the mouse obtained rewards in ALT1. However, this phasic DA releases became weaker in ALT2, then eventually dropped below zero in ALT3 (“DA dip,” see also **Figure 7**). The correlation between the trial order and reward responses was significantly negative in ALT1-3 (correlation coefficient r_rwd_ = -0.45, p = 1.4e-10), indicating a learning effect on the reward response of DA release in the DMS (**Figure 6c**). When the same mouse was subsequently subjected to the random-reward condition of classical conditioning, both the cue and reward responses of DA release appeared in the DMS. In examination of the NAc in another mouse, phasic DA responses to cue and reward were observed throughout all the alternate- and random-reward sessions (**Figure 6d**). There were minimal or no significant effects of learning on the cue and reward responses of DA release in this mouse (r_cue_ = 0.14, p = 0.070; r_rwd_ = -0.17, p = 0.024, **Figure 6e**). Population analysis (DMS: seven mice, NAc: six mice) of the correlation coefficients revealed that the r_rwd_ of the DMS was significantly more negative than that of the NAc (median r_rwd_; DMS: -0.28, NAc: -0.14, p = 0.035, Mann–Whitney *U* test, **Figure 6f**), while they did not differ in terms of the r_cue_ (DMS: -0.044, NAc: 0.0092, p = 1). These results suggest that the WM-based reward information may be dynamically processed via DA inputs to the DMS as learning progresses.

**Figure 6.**
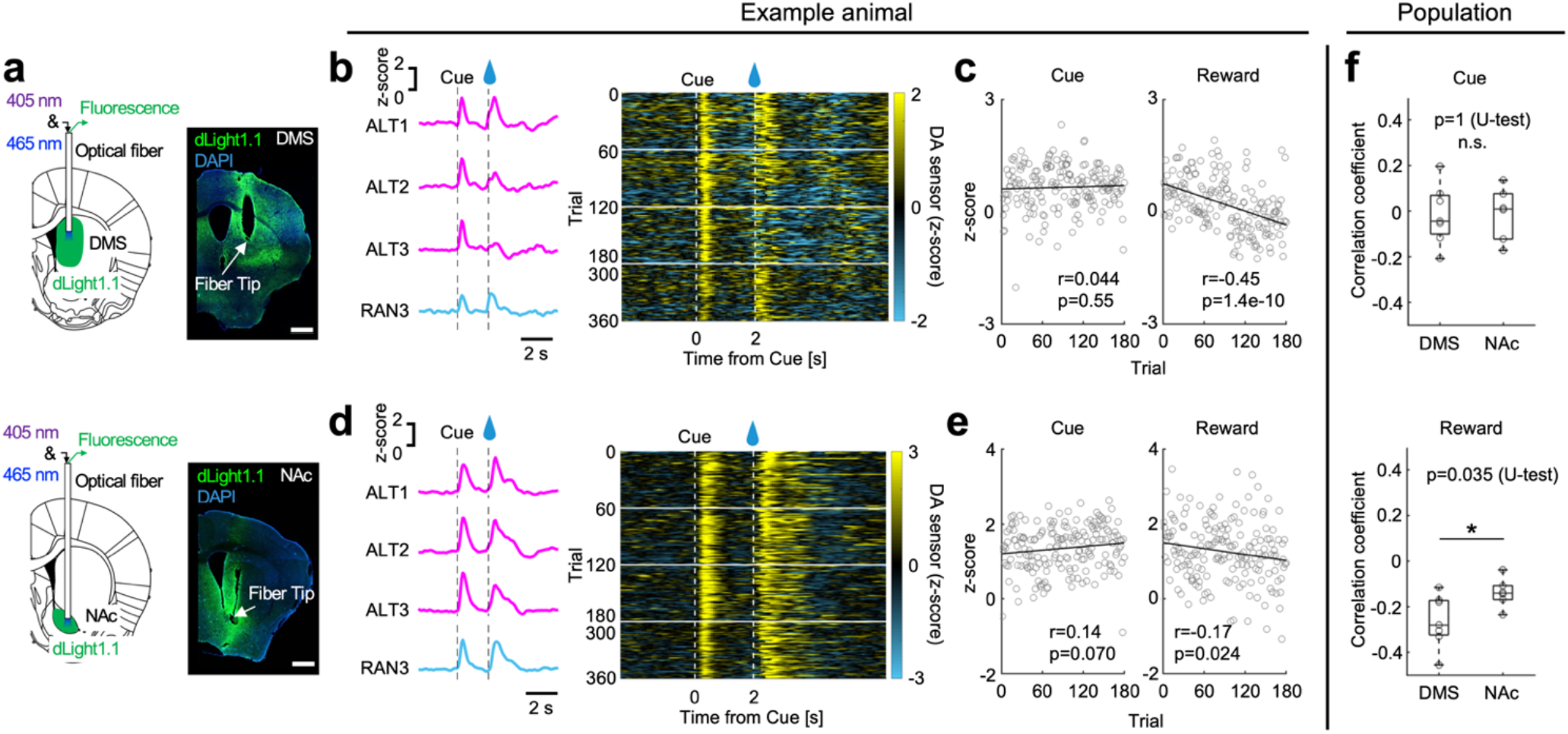
Dopamine release in response to rewards was suppressed in the DMS after sufficient learning of the alternate-reward condition. **(a)** Schematic illustration of the measurement of DA dynamics in the DMS (n = 7 mice) and NAc (n = 6 mice) using fiber photometry, and an example of dLight1.1 fluorescence and optical fiber insertion in the DMS and NAc. Scale bar: 500 µm. **(b)** A representative example of dLight1.1 fluorescence recorded from the DMS. Each line represents the average fluorescence in the reward trials of different task stages, and the heatmap shows trial-by-trial fluorescence. In the alternate-reward condition, the reward response decreased as the task stage progressed. **(c)** Correlation between number of reward trials in the alternate-reward condition and DA release in the DMS. The average fluorescence during 1 s from cue onset and during 1.5 s from reward intake is plotted against the number of reward trials in the alternate-reward condition. DA release in response to rewards significantly decreased with increasing reward trials in the alternate-reward condition, indicating a learning effect. Gray circle and black line indicate averaged fluorescence in each trial and regression line, respectively. Pearson correlation analysis. **(d)** Same as **(b)**, but for the NAc. In the alternate-reward condition, the reward response did not change as the task stage progressed. **(e)** Same as **(c)**, but for the NAc. DA release in response to reward intake was significantly negatively correlated with reward trial experience in the alternate-reward condition. **(f)** Correlation coefficients between the number of reward trials in the alternate-reward condition and DA release. The correlation coefficient between trial experience and DA release in response to the cue stimulus was not significantly different between the DMS and NAc, whereas that between DA release to reward intake was significantly smaller in the DMS than in the NAc, indicating a stronger learning effect in the DMS.

**Figure 7.**
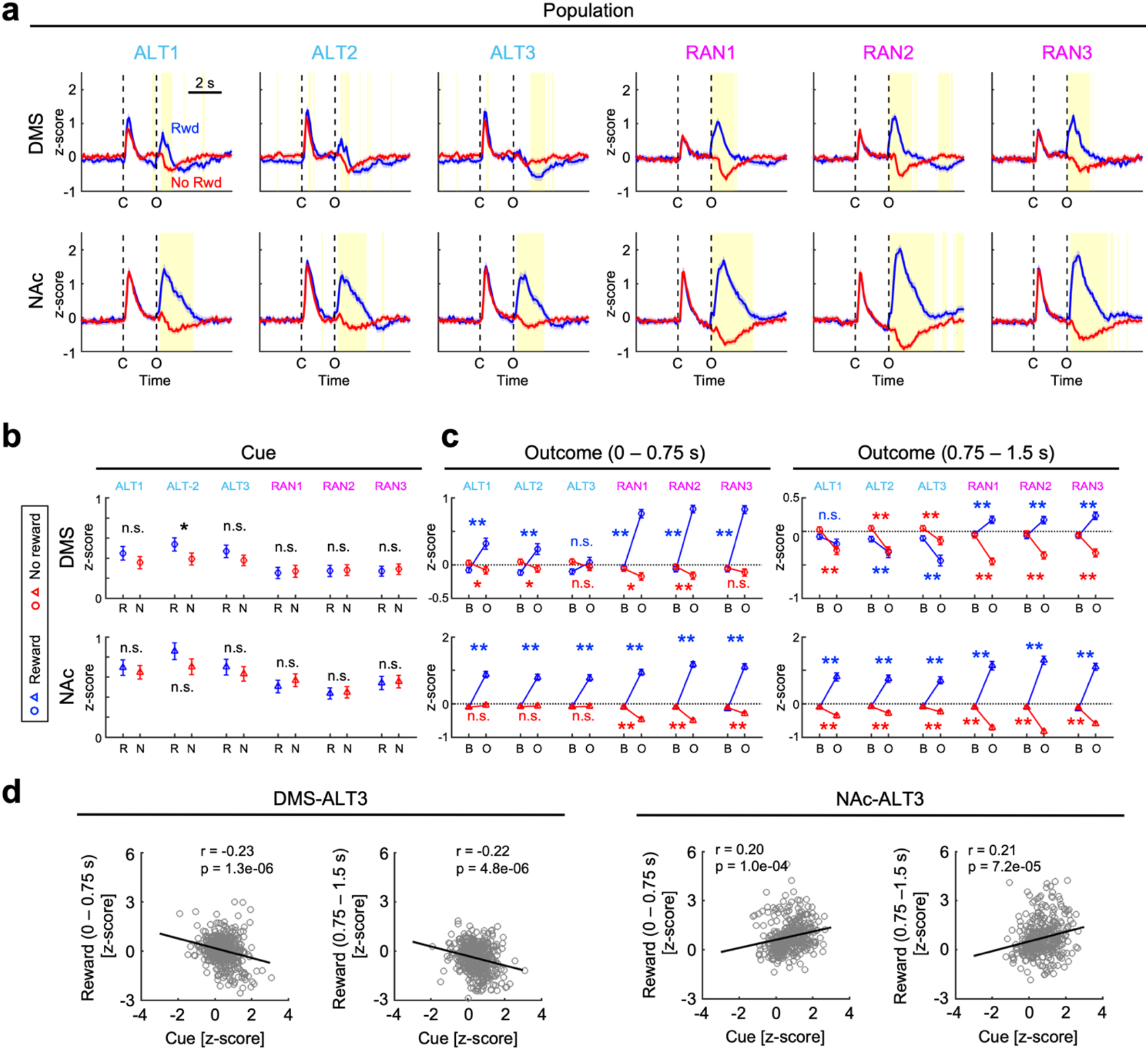
Reward-induced dopamine dips were observed in the DMS after mice learned the reward-alternation rule. **(a)** Population DA dynamics in the DMS (n = 7 mice) and the NAc (n = 6 mice). In the alternate-reward condition, the reward response differed between the DMS and NAc. Blue and red shadows indicate 95% confidence intervals. The yellow bins indicate a significant difference in the z-score between reward and no reward trials (p < 0.01, unpaired *t*-test followed by Bonferroni correction). C: Cue onset, O: outcome onset. **(b)** Quantitative analysis of DA release in response to the cue stimulus. R: reward trial, N: no reward trial. *: p < 0.05, Unpaired *t*-test followed by Bonferroni correction. **(c)** Quantitative analysis of DA release in response to outcome. In the first half of the reward intake period, DA release in the DMS decreased with repetition of the alternate-reward condition and reached baseline levels at ALT3. In the second half of the reward intake period, DA release in the DMS was below baseline from ALT1 to ALT3. **:p < 0.01, *:p < 0.05, paired *t*-test followed by Bonferroni correction. B: baseline (averaged z-score during 2 s before cue onset), O: outcome response. **(d)** Correlation between DA release in response to cue stimulus and the reward intake in ALT3. The correlations are significantly negative and positive in the DMS and NAc, respectively. Black lines indicate regression lines. Pearson correlation analysis.

Some mice that completed the classical conditioning sessions (DMS: six mice, NAc: five mice) were additionally trained to perform an operant conditioning task (Extended Data Fig.6). The reward response of DA release in the DMS was smaller in the alternate-reward condition of the operant learning task than in the random-reward condition, whereas there was no difference between them in the NAc. These results were consistent with the above observation in the classical conditioning task.

According to the reinforcement learning theory, if an actual reward is larger than was predicted just before it was received, the RPE signal of dopaminergic neurons will be changed positively, and *vice versa* (if the reward is smaller, the change will be negative). Then, if mice utilize WM-based reward prediction perfectly in the alternate-reward condition, the RPE change will be zero. Based on this viewpoint, we further evaluated WM-based and WM-free DA dynamics in the DMS and NAc as learning progressed (**Figure 7a**). First of all, the cue response of DA release (1-s duration from cue onset) was not significantly different between the reward and non-reward trials in all sessions and in both areas, except for the DMS in ALT2 (**Figure 7b**; p = 0.016, unpaired *t*-test followed by Bonferroni correction).

Next, we analyzed the DA responses to reward and no reward in the DMS and NAc. The reward response of DA release in the DMS appeared biphasic with positive to negative. The reward response of the early positive phase (0 to 0.75 s after the cue offset) gradually decreased along with the learning progress. In the ALT3, it was no longer significantly different from baseline (p = 0.09, paired *t*-test followed by Bonferroni correction, **Figure 7c**). The reward response of the late negative phase (0.75 to 1.5 s after the cue offset) was also significantly lower than baseline in the ALT2 (p = 1.3e-04) and ALT3 (p = 2.1e-12). Consequently, the reward response in the DMS manifested as a large “dip” in the third session. In contrast, the no-reward response in the DMS appeared as a dip in the ALT1 (p = 0.033) and ALT2 (p = 0.022), as expected, but it disappeared in the ALT3 (p = 0.33). Interestingly, the reward response was significantly smaller than the no-reward response in the third session (p = 6.1e-07, unpaired *t*-test). This gradual inversion of the RPE signal during learning, which is inconsistent with the reinforcement learning theory, has not previously been reported. Thereafter, the positive reward response and the negative no-reward response (dip) reappeared during the random-reward sessions RAN1-3. On the other hand, the reward and no-reward responses in the NAc were always positive and negative (or zero), respectively, corresponding to the typical RPE signals predicted by the standard theory.

We also examined the correlation between the cue and reward responses in the alternate-reward condition in a trial-by-trial manner. The DMS showed negative correlations in ALT3 (cue vs. early reward responses: r = -0.23, p = 1.3e-06, cue vs. late: r = -0.22, p = 4.8e-06, Pearson correlation analysis, **Figure 7d**). In contrast, the NAc showed positive correlations (cue vs. early: r = 0.20, p = 1.0e-04, cue vs. late: r = 0.21, p = 7.2e-05). Note that cue responses in the DMS were not significantly different between reward and no-reward trials in the ALT3 session (p = 0.38, **Figure 7b**). These results suggest that the cue response in the DMS might reflect the degree of confidence for reward prediction in each trial.

## Discussion

In the present study, we investigated how the midbrain DA system contributes to WM-based and WM-free reward prediction by employing ^18^F-FDG-PET imaging, electrophysiological spike recording, and fluorescent fiber photometry in rats and mice seeking alternate (WM-based) or random (WM-free) rewards. Our major findings were as follows. (1) The VTA was differentially activated between the alternate- and random-reward conditions in conjunction with distinct brain areas (**Figures 1 and 2**). (2) Compared to the medial VTA, the lateral VTA represented smaller RPE, reflecting WM-based reward prediction, in the alternate-reward condition (**Figures 3 and 4**). (3) As learning progressed, phasic DA releases in response to alternate rewards and no rewards dynamically changed in the DMS receiving lateral VTA inputs (**Figures 5-7**). Contrary to the reinforcement learning theory, receiving a reward rather than no reward caused a “dip” of DA release in the DMS once the reward alternation pattern was learned well.

### Behaviors reflecting WM-based reward prediction

Typical behavioral learning tasks, whether classical or operant conditioning, require animals to associate different sensory cues or actions with the amount (Cohen et al., 2012; Yoshizawa et al., 2018) or probability (Oyama et al., 2010) of a reward. In contrast, our behavioral tasks always used the same cue and action in all trials in each session, with the only difference being the presence or absence of a reward. The animals did not need to distinguish multiple cues or actions in order to receive a reward. In addition, they were not forced to distinguish between reward and no reward, but naturally learned to predict them. Their reward prediction would manifest just in the reaction time (**Figure 1**) and neural activity in the alternate-reward condition. This simplicity allowed areal comparisons between task sessions for PET imaging and fiber photometry, and neuronal comparisons between trial blocks for electrophysiology. For this natural reward expectation, the animals only needed to operate the WM system (Baddeley, 2003) to maintain one-trial information on the reward in the latest trial (namely, WM-based reward prediction). Thus, our behavioral tasks with reward alternation effectively assess WM-based and WM-free reward prediction in the midbrain.

### Brain areas responsible for WM-based reward prediction

We used ^18^F-FDG-PET imaging, which regards glucose consumption as an indicator of net neural activation (Sokoloff et al., 1977; Phelps et al., 1979), to objectively explore active areas of the whole brain that were related to WM-based reward prediction (Endepols et al., 2010). We found that the Au1, RtTg, PnO, PnC, and VTA activated more strongly in the alternate-reward condition than in the no-reward condition (**Figure 1**). The Au1 might process auditory information on the go-cue tone with more attention or motivation when the reward can be predicted. The RtTg, PnO and PnC are subnuclei of the pontine reticular formation. With regard to auditory processing, the RtTg and PnC are implicated in the acoustic startle reflex (Yeomans and Frankland, 1995; Lee et al., 1996; Koch, 1999; Guo et al., 2021). The RtTg is also involved in motor functions such as eye movements (Gamlin and Clarke, 1995), forelimb movements (Zangger and Schultz, 1978; Matsunami, 1987), and locomotion (Brudzyński and Mogenson, 1984). Hence, reward prediction may allow these areas to reduce the reaction time between the cue and any subsequent action.

The VTA is known as a major source of dopaminergic neurons that encode RPE, which is the discrepancy between actual reward and predicted reward (Schultz et al., 1997; Schultz, 2016). On the whole, the VTA was more activated in the alternate-reward condition than in the random-reward condition. Interestingly, the functional connectivity of the VTA with rostral brain areas was stronger in the alternate-reward condition, while with caudal brain areas it was more substantial in the random-reward condition (**Figure 2**). This suggests that neural circuits between the VTA and rostral brain areas, including the NAc and OFC, may play specific roles in WM-based and WM-free reward prediction. The NAc and OFC receive direct DA inputs from the VTA. In fact, NAc neurons encode state and action value information expected from the environment and from action options (Ito and Doya, 2009, 2015), respectively, and DA release in this area is associated with motivation (Ikemoto and Panksepp, 1999; Mohebi et al., 2019). These studies support our VOI analysis of the VTA and NAc (**Figure 2d**), as the RPE activity would be larger in the VTA when the reward expectation or motivation is lower in the NAc. The OFC neurons also process reward-related information on task structure (Takahashi et al., 2011) and reward uncertainty (Ogawa et al., 2013). Thus, our whole-brain PET analysis shows that the VTA plays a central role in WM-based reward prediction.

### Lateral VTA neurons are involved in WM-based reward prediction

We compared the alternate-, random-, and 100%-reward conditions in terms of the spike activity of single VTA neurons in response to rewards or no rewards in the same rats (**Figure 3**). A population of VTA neurons encoded typical RPE signal, consistent with a previous report (Oyama et al., 2010). According to the reinforcement learning theory, if the rats used WM-based reward prediction, the RPE amplitude with the alternate rewards should be smaller than that with the random rewards. Indeed, relatively smaller RPEs were observed in the alternate-reward condition, especially in the lateral VTA and partly including the boundary with the SNc (**Figure 4**). The reduction of RPE activity by reward alternation may seem inconsistent with the fact that this alternation enhanced the PET signal in the VTA (**Figure 2**). The PET signal is proportional to the metabolic rate of glucose, which reflects synaptic activity (Magistretti and Pellerin, 1996) rather than spike activity. A total number of synaptic inputs, both excitatory and inhibitory, to the VTA neurons might be enhanced in the alternate-reward condition.

Anatomically, the lateral VTA/SNc contains a mixture of NAc- and DMS-projecting DA neurons, while the medial VTA contains mostly NAc-projecting DA neurons (Parker et al., 2016; Saunders et al., 2018). Functionally, NAc-and DMS-projecting neurons preferentially encode reward-related and choice-related information, respectively (Parker et al., 2016). Here we showed that lateral VTA/SNc neurons processed WM-based RPE information, while the medial VTA neurons processed WM-free RPE information. These topological differences presumably extend to the entire cortico-basal ganglia loop. The medial prefrontal cortex (Voorn et al., 2004) engages in WM function (Liu et al., 2014; Bolkan et al., 2017) and may cooperate with the lateral VTA/SNc and DMS in WM-based reward prediction.

### Striatal DA dynamics in WM-based reward prediction during learning

The reinforcement learning theory states that a positive RPE occurs if an actual reward exceeds the value of the expected reward, and a negative RPE occurs if the opposite is true (Schultz et al., 1997; Sutton and Barto, 1998). However, our striatal DA measurements in the DMS showed a negative response (dip) to the actual reward in the alternate-reward condition after the establishment of classical conditioning (**Figures 5-7**). Here, we propose a hypothesis that explains this unexpected phenomenon regarding WM-based and RM-based reward predictions (**Figure 8**). It is assumed that the rats used V_WM_, defined as WM-retained reward expectation based on one-trial information from the latest outcome, and V_RM_, defined as RM-retained reward expectation based on general information from cue stimuli across trials (**Figure 8a**). V_WM_ increases upon no reward and decreases upon reward, so as to anticipate the next outcome. V_RM_ reflects the average reward value in association with the cue stimuli regardless of previous outcomes. V_WM_ and V_RM_ are integrated to calculate the time differentiation, d/dt(V_WM_ + V_RM_).

**Figure 8.**
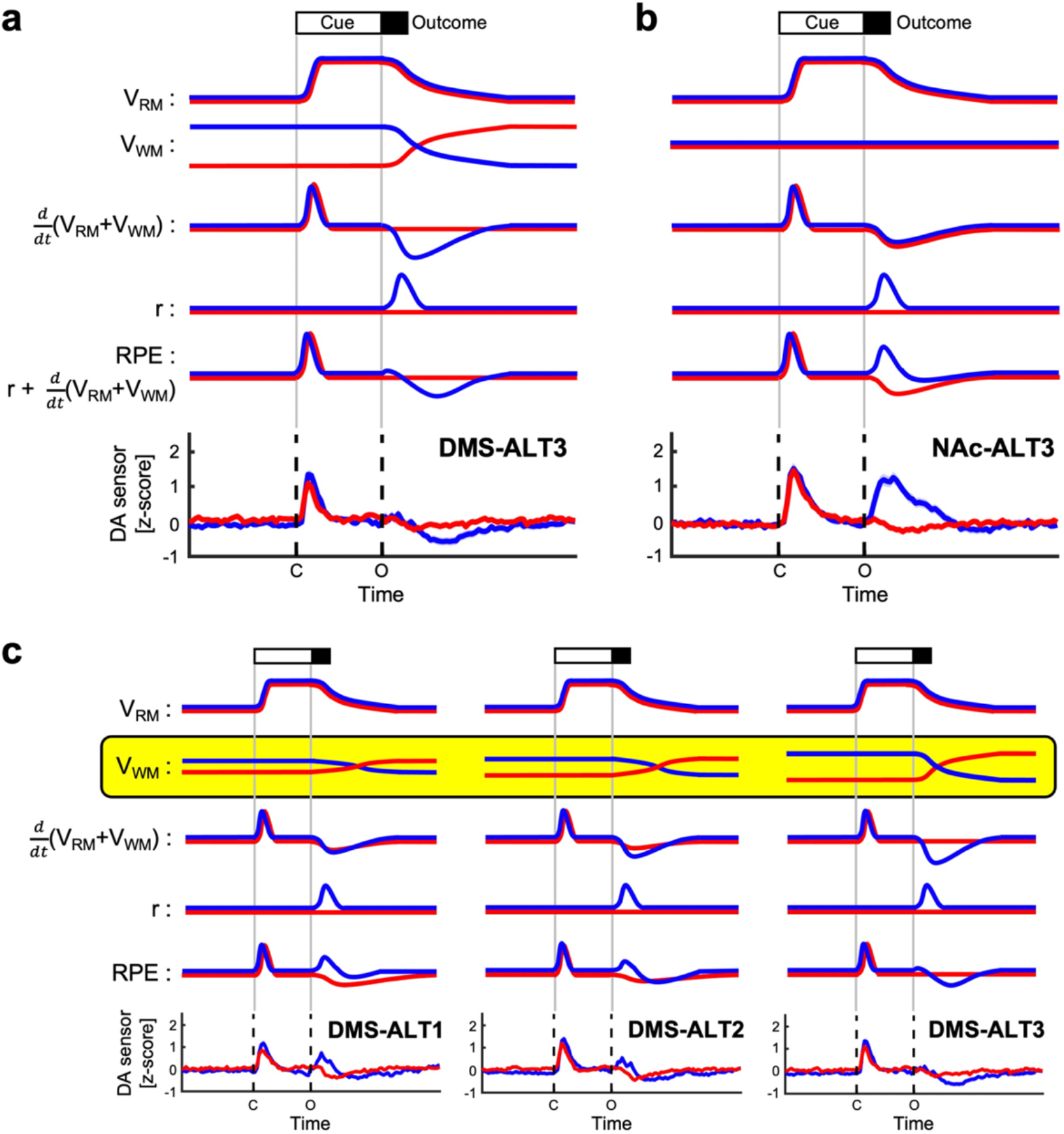
Dopamine release in the DMS reflected the WM-based RPE. **(a)** Schematic model of WM-based reward prediction in the alternate-reward condition. The V_RM_, representing the reward expectation stored in reference memory, rises with cue presentation. The V_WM_, representing the reward expectation stored in working memory, falls at the time of reward intake if reward omission is predicted in the next trial, whereas it rises at the time of reward omission if reward acquisition is predicted in the next trial. Since RPE is the sum of the actual reward *r* and the time change in reward expectation *d/dt*(V_RM_+V_WM_), it is consistent with the DA dynamics observed in the DMS in ALT3. **(b)** Schematic model of WM-free reward prediction under the alternate-reward condition. V_RM_ rises with cue presentation. In contrast to the WM-based reward prediction, V_WM_ assumes a constant value throughout the task period regardless of outcome. In this case, the RPE is consistent with the DA dynamics observed in the NAc in ALT3. **(c)** Schematic model of the formation process of WM-based reward prediction. Assuming that changes in V_WM_ become larger and faster as mice learn the reward-alternation rule, this would explain the changes in DA dynamics in the DMS from ALT1 to ALT3.

Then, we can compute the RPE by summing this predicted reward (d/dt(V_WM_ + V_RM_)) and the actual reward (r). Consequently, in the alternate-reward condition, a positive RPE phase occurs at the time of cue presentation regardless of subsequent outcome, and a negative RPE phase occurs at the time of reward acquisition, thus accurately reproducing the real DA dynamics of the DMS. The biphasic (early-late) property of reward responses (**Figure 7**) results from the different time courses of the predicted and actual rewards. In the absence of V_WM_, the RPE behaves according to the standard theory, which is consistent with the DA release in the NAc (**Figure 8b**). Taken together, our hypothesis supports the idea that the DMS, but not the NAc, processes both WM- and RM-based reward prediction.

Furthermore, our hypothesis successfully explains the development of RPEs in the DMS by assuming that as learning progresses, V_WM_ switches faster and between increasingly different values, with V_RM_ remaining stable (**Figure 8c**). The different time courses of V_WM_ and V_RM_ indicate that WM- and RM-based reward predictions are independently processed in the brain. Our hypothesis also denotes that the RPE depends on the balance between V_WM_, V_RM_, and r. In fact, we failed to observe negative (dip) reward responses in operant conditioning, unlike classical conditioning, in the same mice (Extended data Fig.6). The necessity of operant action would probably increase the value of reward r over the time differentiation of V_WM_ and V_RM_ in our operant conditioning task.

Very recently, (Ishino et al., 2022) reported inverse RPE activity (i.e., a positive response to the unexpected lack of a reward) in the anterior VTA, with corresponding DA release in the NAc (as a preprint: https://doi.org/10.21203/rs.3.rs-1391246/v1). However, our findings differ greatly from their interesting observations. First, we detected no inverse RPE neurons in the lateral or medial VTA (**Figure 4**). Second, we identified a DA dip in response to rewards in the DMS, but not in the NAc (**Figure 7**). Third, and most importantly, the DA dip occurred in a WM-based manner with expected (alternate) rewards, but not with unexpected (random) rewards (**Figure 7**). Different VTA functions may be distributed separately along the M-L and A-P axes.

Striatal DA release may be modulated by local presynaptic mechanisms. For example, dynorphin, an endogenous opioid, inhibits striatal DA release by activating kappa-opioid receptors at the axon terminals of midbrain DA neurons (Narita et al., 2005). Acetylcholine can evoke action potentials at axon terminals, leading to locally initiated DA release into the striatum (Liu et al., 2022). Such presynaptic modulations might underlie the discrepancy between the spike activity of VTA neurons and striatal DA release (Mohebi et al., 2019).

The present study demonstrated that lateral VTA neurons encoded an RPE signal in a WM-based manner, and that their DA release in the DMS dynamically changed along with the progress of outcome learning. Patients with psychiatric disorders such as schizophrenia and depression often display impaired WM function (Forbes et al., 2009; Millan et al., 2012; Lever et al., 2015; Snyder et al., 2015). Our findings will provide novel insights into the physiology and pathophysiology of action optimization based on WM-retained information related to actions and outcomes in these circumstances.

## Materials and Methods

### Ethics

All recombinant DNA and animal experiments in this study were approved by the Institutional Animal Care and Use Committee of Hokkaido University (protocol #17-0045, #22-0023), by the Institutional Animal Care and Use Committee of Tokyo Medical and Dental University (protocol #A2019-274) and by the Institutional Animal Care and Use Committee (IACUC) of RIKEN, Kobe Branch (protocol #MA2006-07).

### Subjects of PET experiments

Male Long-Evans rats (n = 59, 224–280 g body weight, 10 weeks old at the first PET imaging session) were housed individually under a light/dark cycle (lights on at 7:00, off at 19:00). Experiments were performed during the light phase. Water was restricted to 2–4 ml/d during the experimental period. Food was provided *ad libitum* for the entire period.

### Behavioral tasks in PET experiments

Freely moving rats were trained to perform a lever-push task to obtain a water reward. All training and recording procedures were conducted in a 32 × 21.5 × 15 cm custom-built experimental chamber placed in a sound-attenuating box. The chamber was equipped with a spout-lever in front of a small window (3.5 × 3.0 cm) on one wall. A computer program written in Python was used to control a speaker and water pump, and to monitor the states of the lever. Each trial began with a houselight on. When the rat pushed the lever at its own pace and held the push for 500 ms during the houselight on, a go-cue tone (frequency: 3.6 kHz, duration: 100 ms) was presented. When the rat released the lever after the go-cue onset, either a reward (0.1% saccharin water; 10 µL) or no reward (0 µL) was presented, followed by the houselight off and an ITI. We designed two reward presentation conditions. In the alternate-reward condition, reward and no-reward trials were presented alternately, whereas in the random-reward condition, they were presented randomly with equal probability. The ITI was set depending on the number of trials in the previous session in order to align the number of trials between the two conditions.

### PET scanning

Before [^18^F]FDG-PET scanning, all rats were trained on one of these conditions for at least 2–3 days. On the day of PET scanning, each rat received tail vein cannulation at least 1 h prior to the scan while under anesthesia with a mixture of 1.5% isoflurane and nitrous oxide/oxygen (7:3). Following complete recovery, rats performed the experiment with the alternate- or no-reward condition in the operant chamber (alternate: 30 rats, random: 29 rats). After 10 min, rats received an intravenous injection of [^18^F]FDG (ca. 75 MBq/0.4 mL) under freely moving conditions and continued to perform the task for 30 min thereafter. After a 45-min uptake period, rats were anesthetized with a mixture of 1.5% isoflurane and nitrous oxide/oxygen (7:3) and were placed in the gantry of a PET scanner (microPET Focus220, Siemens Co., Ltd, Knoxville, TN, USA). Fifty-five minutes after the [^18^F]FDG injection, a 30-min emission scan was performed. During the PET scan, body temperature was kept at approximately 37℃ with a heating blanket. Emission data were acquired in list mode, sorted into a single sinogram, reconstructed by standard 2D filtered back projection (FBP) with a ramp filter and a cutoff frequency of 0.5 cycles/pixel, or by a statistical maximum a posteriori probability algorithm (MAP) with 12 iterations and point spread function effect.

### Image analysis

Each MAP-reconstructed FDG image was co-registered to an FDG image template using a mutual information algorithm with Powell’s convergence optimization method provided by the PMOD software package (ver. 3.6, PMOD Technologies, Ltd., Zurich, Switzerland). Then the FDG template image was transformed into an MRI reference template that was placed in the Paxinos and Watson stereotactic space (Paxions and Watson, 1998). The transformation parameters estimated from individual MAP-reconstructed FDG images were applied to each FBP-reconstructed FDG image. Subsequently, the voxel size was resampled at 0.12 × 0.12 × 0.12 mm. To enhance the statistical power, each FBP image was spatially smoothed with an isotropic Gaussian kernel (0.6-mm full width at half maximum).

Voxel-based statistical analysis was performed using SPM8 software (Wellcome Department of Imaging Neuroscience, London, UK). Proportional scaling was used for global normalization. A two-sample *t*-test was used for calculating the statistical differences between groups. Functional connectivity was subsequently estimated based on VTA regional activity in which the mean value of the FDG uptake in the VTA at each session was used as a covariate to find regions showing significant correlation across scans. Fisher’s Z-transformation test was applied to assess the significance of difference between two correlation coefficients. The statistical threshold was set at p < 0.0125 (uncorrected) with an extent threshold of 50 contiguous voxels for simple *t*-test analysis, and p < 0.005 (uncorrected) with an extent threshold of 50 contiguous voxels for functional connectivity analysis.

### Subjects of electrophysiological recording

Male Long-Evans rats (n = 4, 220–283 g body weight, 8 weeks old at surgery) were housed individually under a light/dark cycle (lights on at 9:00, off at 21:00). Experiments were performed during the light phase. Water was restricted to 5–8 ml/d during the experimental period. When necessary, an agar block (containing 15 ml water) was given to the rats in their home cage to maintain >85% of their original body weight (Soma et al., 2017). Food was provided *ad libitum* for the entire period.

### Surgery for rat electrophysiological recordings

Rats were placed in a stereotaxic frame (SR-10R-HT, Narishige, Tokyo, Japan) and anesthetized with isoflurane (4.5% for induction and 2.0–2.5% for maintenance; body temperature 37 °C). Reference and ground electrodes were implanted above the cerebellum, a head plate (CFR-2, Narishige) was attached to the skull using small anchor screws and pink dental cement (Unifast 2, GC, Tokyo, Japan) verified for brain effects (Yoshizawa and Funahashi, 2020). The exposed surface of the skull and brain was covered with silicone sealant (Dent Silicone-V, Shofu, Kyoto, Japan). Analgesics and antibiotics were applied postoperatively as required (meloxicam, 1 mg/kg s.c.; 0.1% gentamicin ointment, *ad usum externum*).

### Behavioral task in electrophysiological recording

One week after recovering from surgery, rats were head-fixed using the head plate and were habituated to a restraint operant chamber (TaskForcer, O’Hara, Tokyo, Japan) for 1–2 d before task training. They spontaneously started each trial by pushing a spout-lever in the chamber with their right forelimbs and holding it for a short period (750–1250 ms, **Figure 3b**). After the holding period, a go-cue sound (frequency: 4 kHz, duration: 100 ms) was presented to instruct them to pull the spout-lever. When they did so, a success-cue sound (frequency: 10 kHz, duration: 500 ms) was presented, followed by a delay (800–1200 ms) and an outcome. In the alternate-reward condition, a drop of 0.1% saccharin water (10 µL) was alternately delivered or not delivered, whereas in the random-reward condition, it was randomly delivered with 50% probability. In the 100%-reward condition, it was always delivered. The alternate- and random-reward conditions were switched every 50–70 trials (**Figure 3c**). The 100%-reward condition consisted of 150 trials and appeared every 10 times the alternate- and random-reward conditions were switched. The alternate-, random- and 100% reward conditions were switched without any external cue.

### Electrophysiological recording

Once the rats completed training of the behavioral task, they underwent a second surgery in which a tiny hole was made in the skull and an electrode was inserted into the brain under anesthesia. Extracellular multichannel recordings were performed using a 32-channel silicon probe (a32-Isomura-6-14-r2-A32 or ISO-3x-tet-A32, NeuroNexus Technologies, Ann Arbor, MI, USA) from the left VTA (A-P: -5.2 ∼ - 5.4, M-L: 0.55 ∼ 1.4, DV: 7.8 ∼ 8.2 from the brain surface) under head fixation without anesthesia. The probe was inserted into and removed from the VTA in every daily session. The exposed surface of the skull and brain was covered with silicone sealant (Dent Silicone-V, Shofu) after completing each recording session. The track of the silicon probe was histologically confirmed later by fluorescence microscopy (**Figure 4c**).

The extracellular signals were amplified (final gain ×2000) and filtered (0.5 Hz to 10 kHz) through a 32-channel head-stage (MPA32I, Multi-Channel Systems, Reutlingen, Germany) and main amplifier (FA64I, Multi-Channel Systems). The signals were digitized at 20 kHz and 12 bits and recorded with a 32-channel hard-disc recorder (LX-120, TEAC, Tokyo, Japan). These signals included the spike activity of multiple neurons and local field potentials. The raw signal data were processed offline to isolate spike events of individual neurons in each tetrode of the silicon probe, using the semi-automatic spike-clustering software EToS (Takekawa et al., 2010, 2012) and the manual clustering software Klusters, in conjunction with NeuroScope (Hazan et al., 2006). To classify putative VTA dopaminergic neurons and GABAergic interneurons, the spike width of isolated neurons was defined as the time from the first to the second positive peak of the averaged spike waveform (**Figure 4d**). VTA neurons with a spike width <0.4 ms were classified as non-dopaminergic cells.

### Subjects of fiber photometry recording

Male C57BL/6 mice (n = 13; 20–22 g body weight, 8 weeks old at surgery) were housed individually under a 12/12 h light/dark cycle (lights on at 7:00, off at 19:00). Experiments were performed during the light phase. Water was restricted to 1–2 ml/d during the experimental period. When necessary, an agar block (containing 1 ml water) was given to the mice in their home cage to maintain >85% of their original body weight (Yoshizawa et al., 2018). Food was provided *ad libitum* for the entire period.

### Surgery for fiber photometry recording

Mice were anesthetized with isoflurane (4.0% for induction and 1.0–3.0% for maintenance; body temperature 37 °C) and placed in a stereotaxic frame. The skull was exposed, a hole (diameter: 1.0 mm) was drilled in the skull, and the dura was removed over the imaging site. For fiber photometry recording, 400 nl of AAV2/5-CAG-dLight1.1 (1.7×10^13^ GC/ml, 111067-AAV5, Addgene, Watertown, MA, USA) was slowly injected into the DMS (n = 7, A-P: +0.5, M-L: +1.75, D-V: 2.85 from the brain surface) or NAc (n = 6, A-P: +1.3, M-L: +1.25, D-V: 4.25 from the brain surface) using a microsyringe pump (Legato100, Kd Scientific, Holliston, MA, USA). After the AAV injection, an optical fiber (diameter: 400 µm, length: 5 mm, MFC_400/430-0.66_5mm_ZF1.25(G)_FLT, Doric, Quebec, Canada) was implanted 200 µm above the AAV injection coordinates. The optical fiber was fixed with UV adhesive (Loctite 4305, Henkel, Düsseldorf, Germany) and clear dental cement (Super bond, Sun Medical, Shiga, Japan). A head plate (CF-10, Narishige) was fixed with pink dental cement (Unifast 2, GC). Analgesics and antibiotics were applied postoperatively as required (meloxicam, 1 mg/kg s.c.; 0.1% gentamicin ointment, *ad usum externum*).

### Tone-reward association task for fiber photometry recording

Three weeks after AAV injection and optical fiber implantation, mice were head-fixed using the head plate and habituated to a custom-built restraint operant chamber (Yoshizawa et al., 2018) (O’Hara) for 3–5 d before task training (**Figure 5a**). In training sessions, mice were always rewarded with a drop of 0.1% saccharin water (4 µl) immediately after a tone cue (10 kHz, 2 s). Fiber photometry recordings were started after mice, predicting a reward, licked a water-spout placed in front of their mouths during the cue presentation. In recording sessions, mice performed the task under the alternate-reward condition in which a reward trial and a no-reward trial alternately appeared (**Figure 5b**). If mice licked the spout in the 0.5 s before outcome presentation significantly more frequently in post-no-reward trials than in post-reward trials, the session was deemed successful (p < 0.05, unpaired *t*-test). After three successful sessions, mice performed three sessions in the random-reward condition, in which a reward trial and a no-reward trial randomly appeared with 50% probability (**Figure 5c**). To quantify the difference in reward-predictive licking between the post-no-reward and post-reward trials, the effect size *d* was calculated as follows:

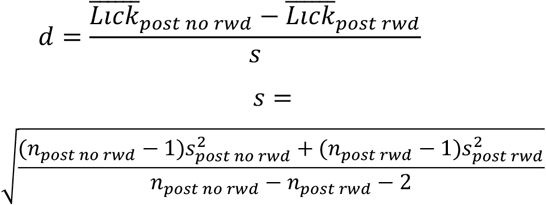

where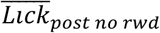 and 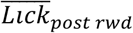 are the average numbers of reward-predictive licks in the 0.5 s before outcome presentation in the post-no-reward and post-reward trials, respectively. 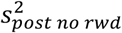 and 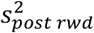 are variances of reward-predictive licking in the post-no-reward and post-reward trials, respectively.*n*_*post no rwd*_ and *n*_*post rwd*_ are the numbers of post-no-reward and post-reward trials, respectively. A daily session consisted of 120 trials. Licks were detected by interruptions of an infrared beam placed in front of the water tube. ITIs were randomly selected from 7 to 13 s.

### Fiber photometry recording

In the head-fixed mice in this study, we used a previously published fiber photometry recording protocol for freely-moving mice (Patel et al., 2020). The fiber photometry system consisted of two excitation channels. A 465 nm LED (CLED_465, Doric) and a 405 nm LED (CLED_405, Doric) were driven by a LED controller (LEDD_4, Doric) to obtain a DA-dependent signal and a DA-independent isosbestic signal, respectively. These LEDs were alternately turned on and off at 13.3 Hz. Fluorescence from dLight and isosbestic fluorescence were directed through dichroic mirrors (iFMC6_IE(400-410)_E1(460-490)_F1(500-540)_E2(555-570)_F2*(580-680)_S, Doric) and were acquired using a photodetector. The signals were passed through a 10x amplifier and were sampled at 1 kHz with a data acquisition system (Power1401, Cambridge Electronic Design, Cambridge, UK). The acquired photometry signals were processed with custom-written code in MATLAB (MATLAB R2018a, Mathworks, Natick, MA, USA). First, the signals were downsampled to 13.3 Hz for further analysis. A fitting curve was estimated and subtracted from the original signal to remove exponential and linear signal decay during the recording session. A linear fit was applied to align the 405-nm signal to the 465-nm signal, and then the fitted 405-nm signal was subtracted from the 465-nm channel and divided by the fitted 405-nm signal to calculate ΔF/F values. The ΔF/F time-series trace was normalized using z-scores to account for data variability across animals and sessions.

### Immunohistochemistry

To confirm the track of the optical fiber, after every fiber photometry recording was completed, mice were deeply anesthetized with pentobarbital sodium and then perfused with 4% PFA. Brains were carefully removed so that optical fibers would not cause tissue damage, post-fixed in 4% PFA at 4 °C overnight, and then transferred to a 30% sucrose / phosphate-buffered saline (PBS) solution at 4 °C until brains sank to the bottom. Coronal sections were cut at 50 µm on an electrofreeze microtome (FX-801, Yamato, Saitama, Japan) and stored in wells containing PBS at 4 °C. Free-floating sections were washed four times in PBS for 15 min and placed in blocking buffer containing 10% normal donkey serum (017-000-121, Jackson ImmunoResearch Laboratories, West Grove, PA, USA) and 0.1% Triton X-100 in PBS for 1 h at room temperature. Sections were simultaneously incubated in primary antibody chicken anti-GFP (GFP-1010, Aves Labs, Davis, CA, USA) diluted 1:500 in blocking buffer overnight at 4 °C. Afterward, sections were washed four times for 15 min in PBS and temporarily stored at 4 °C. Sections were then incubated in secondary antibody donkey anti-chicken Alexa Fluor 488 (703-545-155, Jackson ImmunoResearch Laboratories) diluted 1:500 in blocking buffer overnight at 4 °C. The next morning, sections were washed four times for 15 min in PBS, mounted on glass slides and coverslipped with DAPI-Fluoromount-G (SouthernBiotech, Birmingham, AL, USA). A fluorescence microscope (Eclipse Ci-L, Nikon, Tokyo, Japan) was used to inspect stained tissue and pictures were taken using NIS-Elements software (NIS-Elements D, Nikon).

### Statistical analyses

We used appropriate statistical tests, i.e., paired or unpaired *t*-test, Mann-Whitney *U* test, Wilcoxon signed rank test, and Chi-squared tests with or without Bonferroni’s multiple comparison. Differences were considered statistically significant when p < 0.05. See “Results” for details.

**Extended Data Fig.1.**
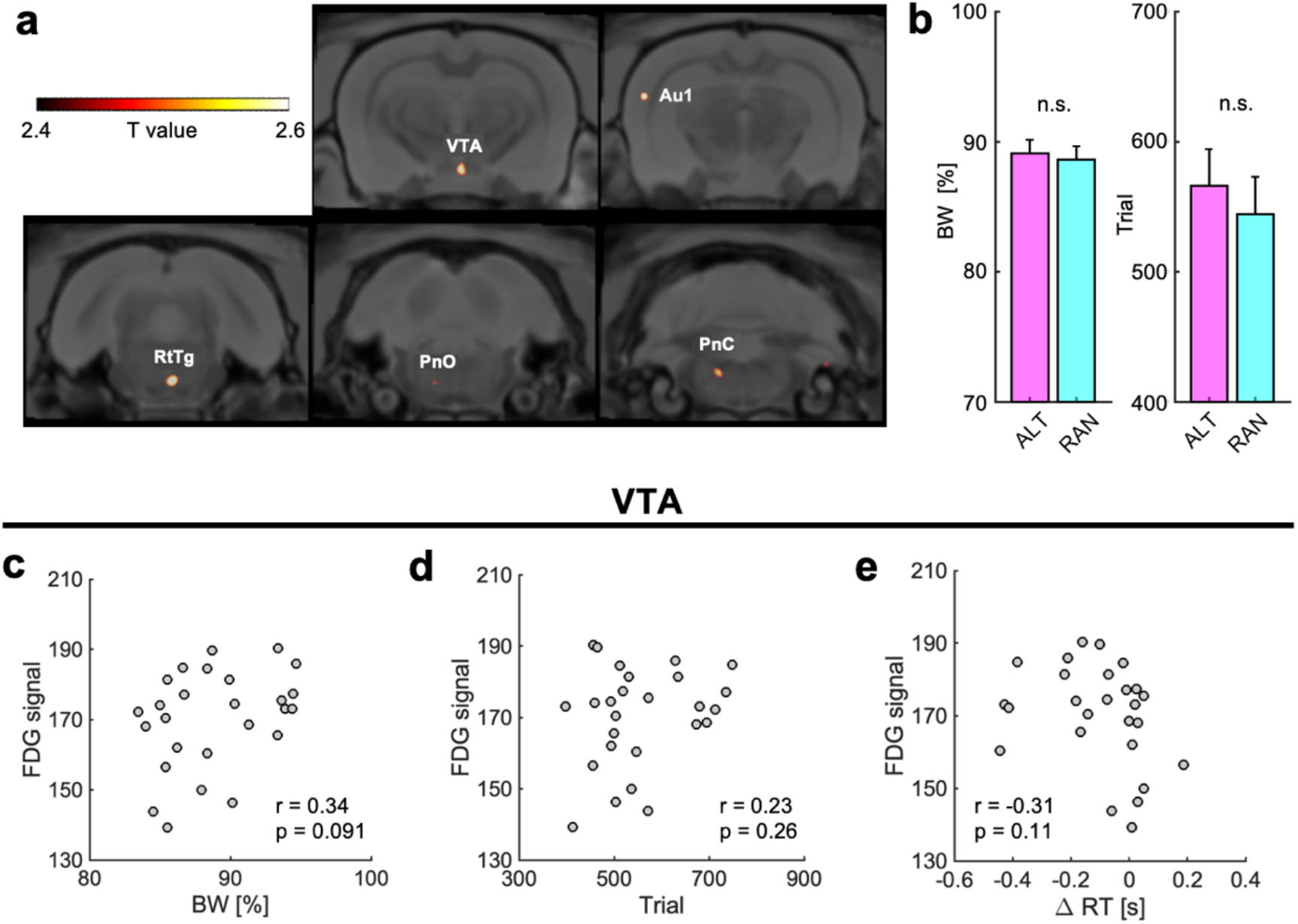
No effect of behavioral parameters on [^18^F]fluorodeoxyglucose signal. **(a)** Coronal views of activated brain regions in the alternate-reward condition. Au1: primary auditory area, VTA: ventral tegmental area, RtTg: reticulotegmental nucleus of the pons, PnO: pontine reticular nucleus, oral part, PnC: pontine reticular nucleus, caudal part. p < 0.0125, uncorrected, height threshold: T = 2.39, n = 13 rats. **(b)** Percentage change of rat body weight (BW) after water restriction and numbers of trial per session were not significantly different between in the alternate- and random-reward conditions. n.s.: p ≥ 0.05, unpaired *t*-test. **(c)** There was no significant correlation between percentage change of BW and [^18^F]fluorodeoxyglucose (^18^F-FDG) signal strength in the VTA. R = 0.34, p = 0.091, Pearson correlation analysis. **(d)** Same as (c), but for number of trials per session. r = 0.23, p = 0.26. **(e)** Same as (c), but for difference in reaction time (RT) between the alternate- and random-reward conditions. r = -0.31, p = 0.11

**Extended Data Fig.2.**
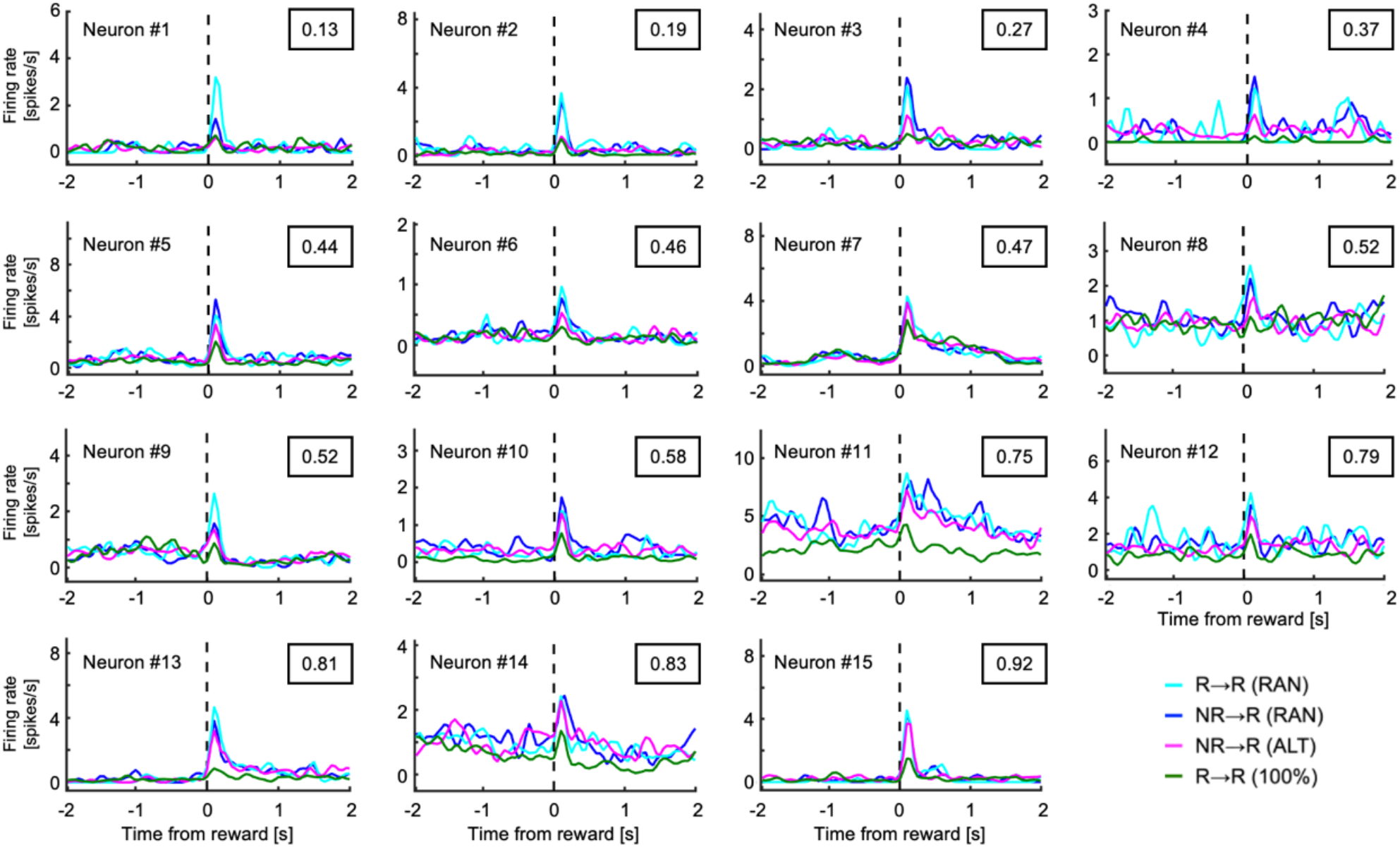
Reward responses of all reward prediction error neurons. Neuron indices sorted on the value of relative amplitude. The value in each square represents the relative amplitude. R: reward trial, NR: no-reward trial.

**Extended Data Fig.3.**
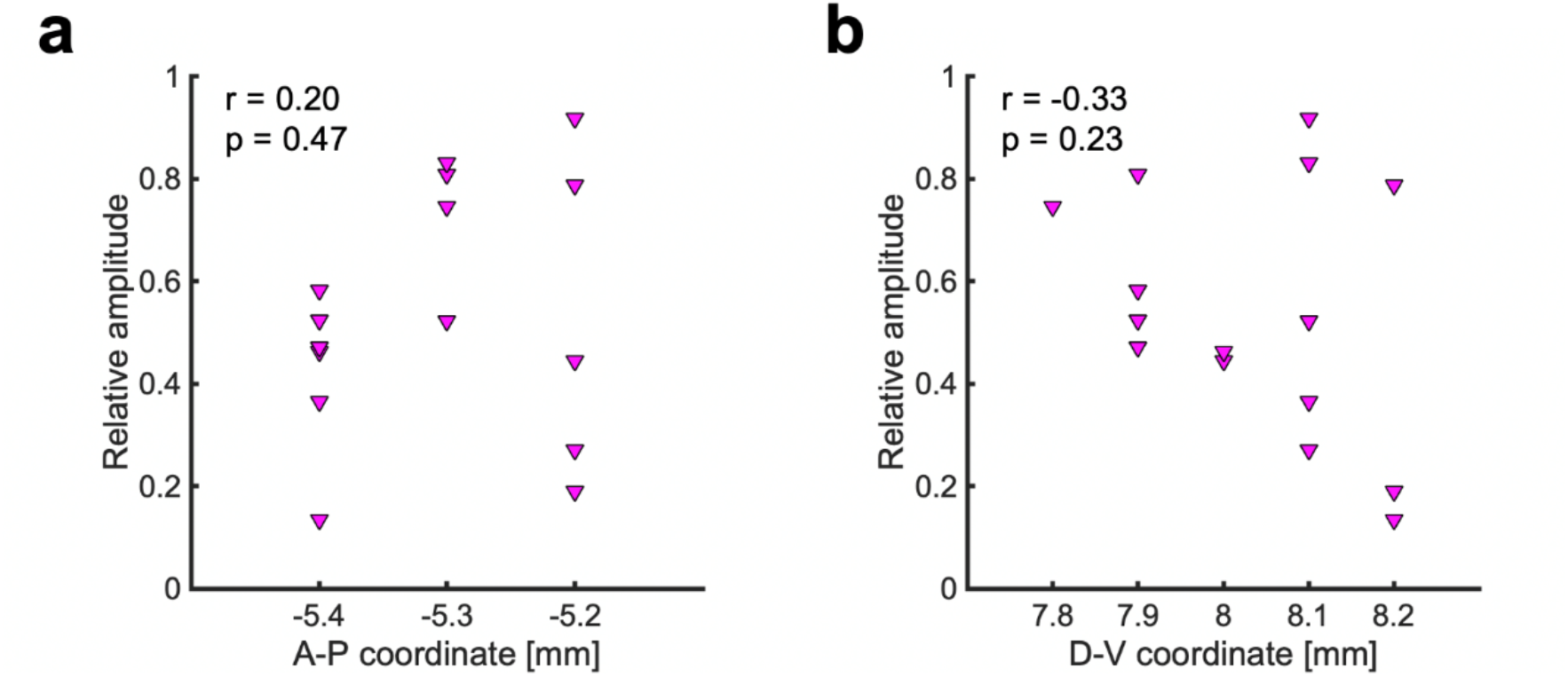
Relation between recording position and response to the alternate reward. **(a)** The relative amplitude of each RPE neuron was not significantly correlated with the anterior-posterior (A-P) coordinate of the recording position. r = 0.20, p = 0.47, Pearson correlation analysis. **(b)** Same as (a), but for the dorsal-ventral (D-V) coordinate. r = -0.33, p = 0.23.

**Extended Data Fig.4.**
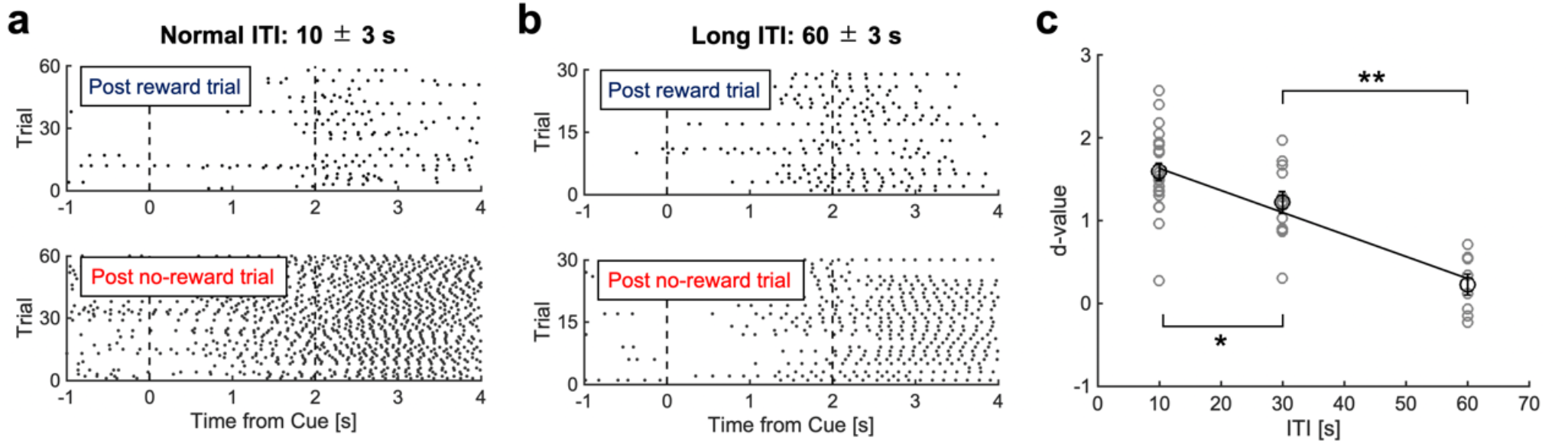
Effects of intertrial interval extension on reward-predictive behavior. **(a)** When the intertrial interval (ITI) was set to 10 ± 3 s in the alternate-reward condition, the licking frequency during the cue period was higher in the post-no-reward trial than in the post-reward trial. In the alternate-reward condition, since the trial after the no-reward trial was a reward trial, mice performed WM-based reward prediction with the outcome information of the previous trial retained in WM. **(b)** When the ITI was set to 60±3 s in the same task, the licking frequency during the cue period was similar between the post no-reward and post-reward trials. ITI prolongation caused the outcome information of the previous trial to disappear from WM, and the RM-based reward prediction arising from the associative learning of cue and reward became apparent. **(c)** In the alternate-reward condition, the ITI was set to 10 ± 3 (Normal ITI), 30 ± 3 (Middle ITI), and 60 ± 3 (Long ITI) s. The difference in the frequency of licking for 0.5 s before the outcome onset was compared between the post-no-reward and post-reward trials using the d-value. The d-values were significantly smaller for Middle ITI than for Normal ITI, and for Long ITI than for Middle ITI. d-values could be regressed from ITI with high accuracy. Black circles and error bars indicate mean ± s.e.m. **: p < 0.01, unpaired *t*-test, n = 4 mice.

**Extended Data Fig.5.**
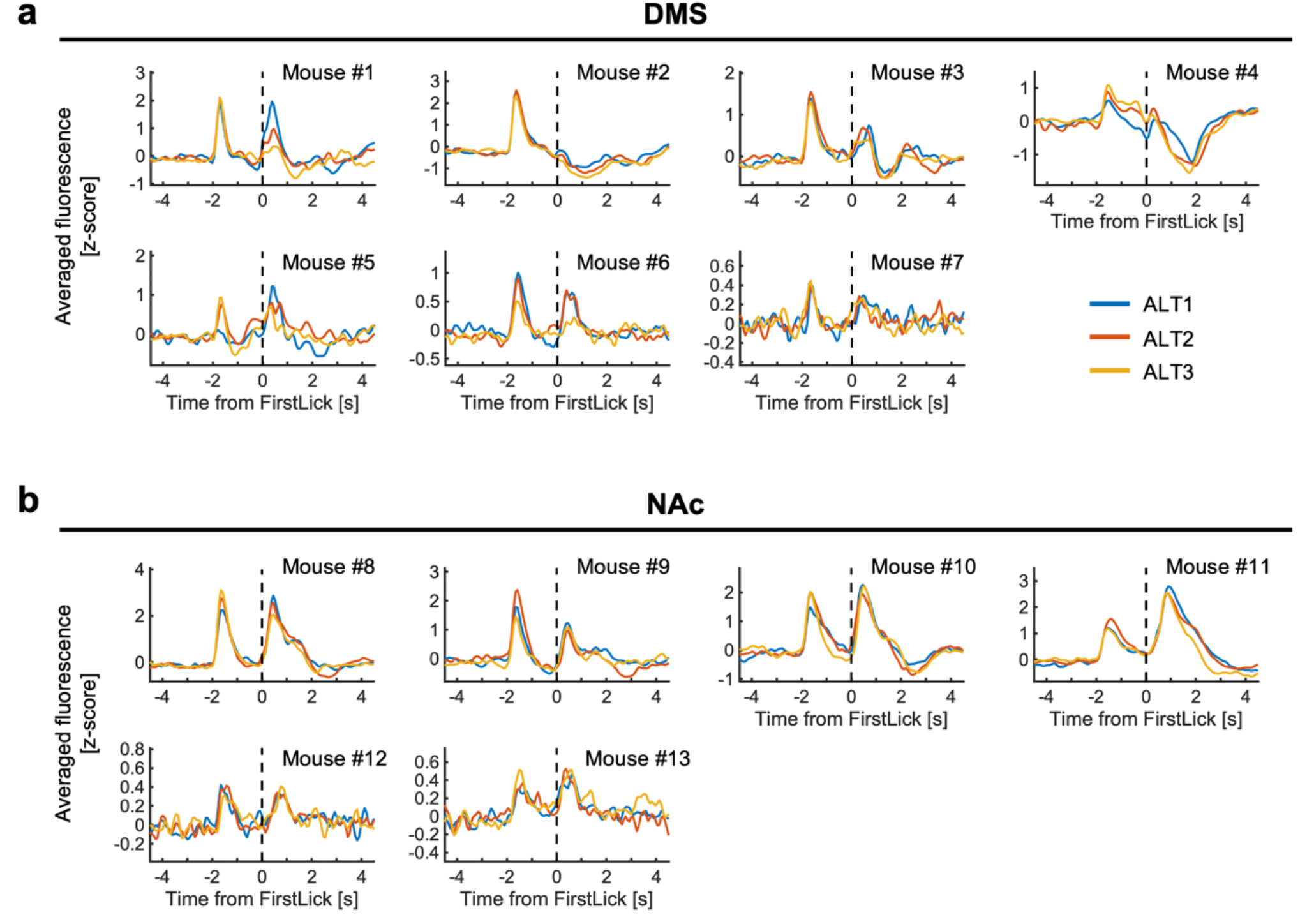
Individual DA dynamics in reward trials in the alternate-reward condition. **(a)** All dLight1.1 fluorescence data recorded from the DMS. Each line represents averaged fluorescence in the reward trials of different task stages in the alternate-reward condition. **(b)** Same as (a), but for the NAc.

**Extended Data Fig.6.**
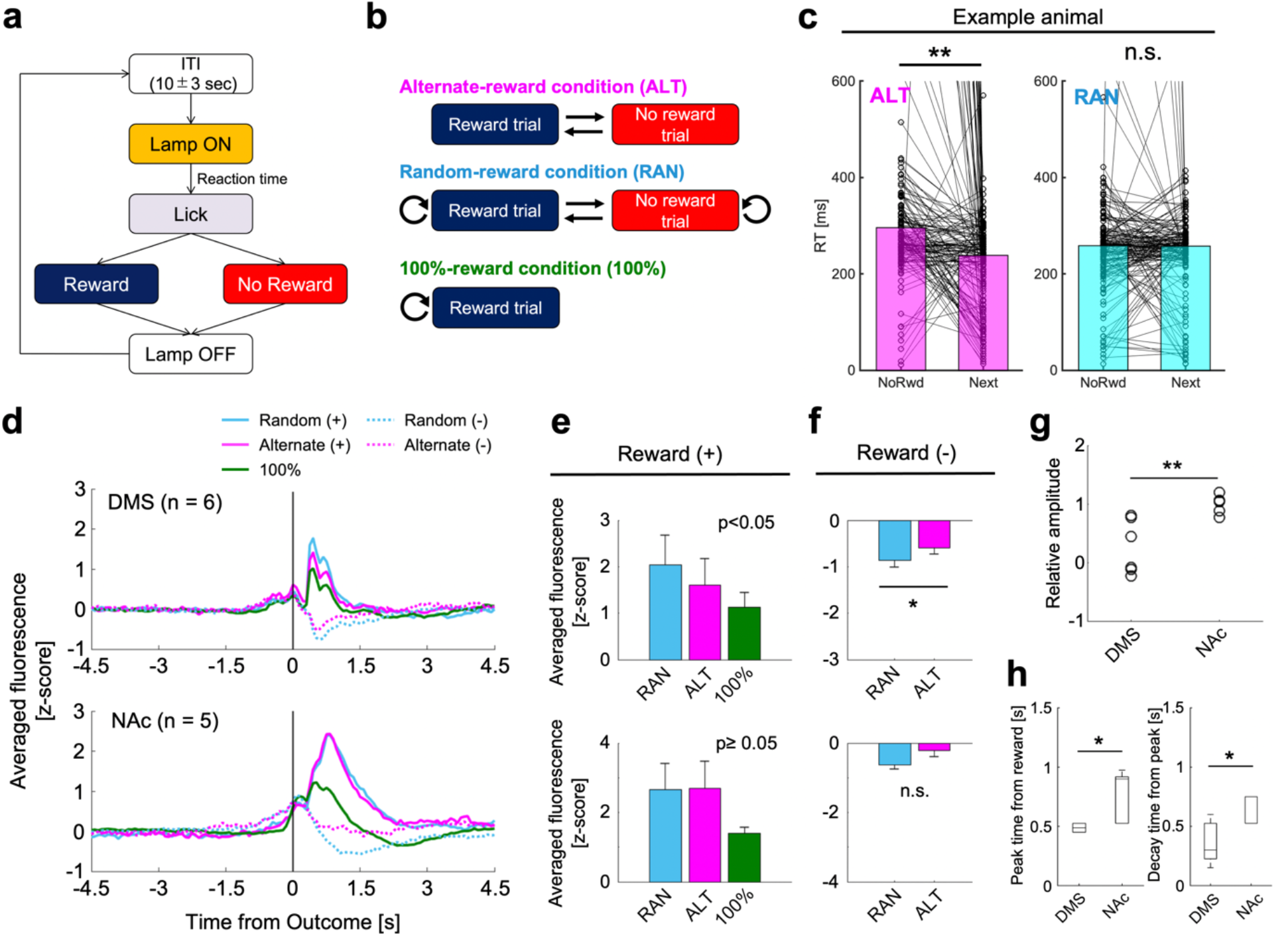
DA dynamics in the DMS and NAc during an operant task. **(a)** Diagram of an operant task for DA fiber photometry. The head and body of each mouse was restrained by a metal frame and tube, as in Figure 5a. A water spout was placed in front of its mouth. Each trial began with turning on a house lamp. When the mouse spontaneously licked the water spout (without any cue signal), a drop of 0.1% saccharin water (4 µl) was or was not presented immediately. At the end of each trial, the house lamp was turned off, followed by a 10±3 s ITI. **(b)** Reward conditions. The alternate- and random-reward conditions were the same as in Figure 5b. In the 100%-reward condition, mice were always rewarded. **(c)** A representative example of RTs in a single session. In the alternate-reward condition, RTs were shorter in the trials following no-reward trials. In the random-reward condition, the previous outcome did not affect the RT in the next trial. **(d)** Population DA dynamics in the DMS (n = 6 mice) and NAc (n = 5). **(e)** Comparison of DA release in the reward trial. In the DMS, DA releases in response to rewards were significantly different between reward conditions (p < 0.05, one-way ANOVA), whereas in the NAc, there was no significant difference (p ≥ 0.05). **(f)** Same as (e), but for the no-reward trial. In the DMS, DA release was more strongly suppressed in the random-reward condition than in the alternate-reward condition, whereas in the NAc, there was no significant difference. *: p < 0.05, n.s.: p ≥ 0.05, unpaired t-test. **(g)** Comparison of relative amplitudes. Relative amplitude was calculated as in Figure 4g, but for dLight1.1 fluorescence. The relative amplitude was significantly smaller in the DMS than in the NAc. **: p < 0.01, unpaired t-test.**(h)** Peak and decay time of DA release. The peak time after reward intake and the decay time after the peak were significantly shorter in the DMS than in the NAc. *: p < 0.05, unpaired t-test.

